# Targeting CD19-positive lymphomas with the antibody-drug conjugate (ADC) loncastuximab tesirine: preclinical evidence as single agent and as combinatorial approach

**DOI:** 10.1101/2023.08.17.553668

**Authors:** Chiara Tarantelli, David Wald, Nicolas Munz, Filippo Spriano, Alessio Bruscaggin, Eleonora Cannas, Luciano Cascione, Eugenio Gaudio, Alberto J. Arribas, Shivaprasad Manjappa, Gaetanina Golino, Lorenzo Scalise, Emanuele Zucca, Anastasios Stathis, Patrick H. van Berkel, Davide Rossi, Paolo F. Caimi, Francesca Zammarchi, Francesco Bertoni

**Author notes:** Co-corresponding authors: Dr Chiara Tarantelli, Institute of Oncology Research, via Francesco Chiesa 5, 6500 Bellinzona, Switzerland., Prof. Francesco Bertoni, Institute of Oncology Research, via Francesco Chiesa 5, 6500 Bellinzona, Switzerland. first co-authors (Chiara Tarantelli, David Wald).

## Abstract

**Purpose:** Antibody-drug conjugates (ADCs) represent one of the most successful therapeutic approaches introduced in clinical practice in the last years. Loncastuximab tesirine (ADCT-402) is a CD19 targeting ADC, in which the antibody is conjugated through a protease cleavable dipeptide linker to a pyrrolobenzodiazepine (PBD) dimer warhead (SG3199). Based on the results of a phase 2 study, loncastuximab tesirine was recently approved for adult patients with relapsed/refractory large B-cell lymphoma.

**Experimental Design:** We assessed the activity of loncastuximab tesirine in *in vitro* and *in vivo* models of lymphomas, correlated its activity with CD19 expression levels and identified combination partners providing synergy with loncastuximab tesirine.

**Results:** Loncastuximab tesirine was tested across 60 lymphoma cell lines. Loncastuximab tesirine has strong cytotoxic activity in B-cell lymphoma cell lines and the *in vitro* activity is correlated with CD19 expression level and with intrinsic sensitivity of cell lines to the ADC’s warhead.

Loncastuximab tesirine was more potent than other anti-CD19 ADCs (coltuximab ravtansine, huB4-DGN462), albeit the pattern of activity across cell lines was correlated. Loncastuximab tesirine activity also largely correlated with cell line sensitivity to R-CHOP.

Combinatorial in vitro and in vivo experiments identified the benefit of adding loncastuximab tesirine to other agents, especially BCL2 and PI3K inhibitors.

**Conclusions:** Our data support the further development of loncastuximab tesirine as single agent and in combination for patients affected by mature B-cell neoplasms. The results also highlight the importance of CD19 expression, and the existence of lymphoma populations characterized by resistance to multiple therapies.

---

Despite the clear improvements achieved in the cure of patients affected by lymphoid neoplasms, novel therapeutic strategies are still needed since current therapies are not yet curative for too many persons (1-3). Antibody-drug conjugates (ADCs) represent one of the most successful therapeutic approaches introduced in clinical practice in the last 25 years (4-6). ADCs are complex compounds that contain three components: an antibody, a warhead (i.e., a cytotoxic agent) and a linker that joins the two together. ADCs enable targeted delivery of potent warheads into tumor cell using antibodies specific for tumor antigens.

Due to its pattern of expression and its biologic role in lymphocytes, the B cell marker CD19 has been heavily exploited for antibody-based therapies, including ADCs, and, more recently, for cellular therapies (5,7-14). Loncastuximab tesirine (ADCT-402) is a CD19 targeting ADC, in which the CD19-specific antibody is stochastically conjugated through a protease cleavable dipeptide linker to a pyrrolobenzodiazepine (PBD) dimer warhead (SG3199) (10). Following binding to CD19 positive cells, loncastuximab tesirine is rapidly internalized and transported to lysosomes where the linker is cleaved to release the PBD dimer SG3199 (10). In contrast with the microtubule-disrupting monomethyl auristatin E (MMAE) used in the CD30-targeting brentuximab vedotin and the CD79B-targeting polatuzumab vedotin ADCs (15,16), SG3199 belongs to a new generation of DNA cross-linking agents. SG3199 binds to guanine residues in the DNA minor groove forming covalent cross-links of the two DNA strands (17,18). Loncastuximab tesirine has been studied in various clinical trials (19-21) and, based on the results of a phase 2 study (19,22), it was recently approved in the USA and Europe for adult patients with relapsed/refractory (R/R) large B-cell lymphoma after at least two prior lines of systemic therapy (23).

Here, we assessed the anti-tumor activity of loncastuximab tesirine in a large panel of lymphoma cell lines, with a focus on the expression of its target and on the identification of active combination partners.

## Materials and Methods

### Cell lines

Lymphoma cell lines were cultured according to the recommended conditions, as previously described (24). All media were supplemented with fetal bovine serum (10% or 20%) and penicillin-streptomycin-neomycin (≈5,000 units penicillin, 5 mg streptomycin, and 10 mg neomycin/mL; Sigma). Human cell line identities were confirmed by short tandem repeat DNA fingerprinting using the Promega GenePrint 10 System kit (B9510). Cells were periodically tested for mycoplasma negativity using the MycoAlert Mycoplasma Detection Kit (Lonza).

### Compounds

Loncastuximab tesirine, SG3199 and B12-SG3249 were provided by ADC Therapeutics. Copanlisib was purchased from MedKoo Biosciences Inc. (Morrisville, NC, USA). Idelalisib, venetoclax, bendamustine, olaparib, ibrutinib, doxorubicin, vincristine, prednisolone, bortezomib, and lenalidomide were purchased from Selleckchem (Houston, TX, USA). Rituximab was purchased from Roche (Basel, Switzerland), and 4-hydroperoxy-cyclophosphamide from Santa Cruz Biotechnology (Heidelberg, Germany).

### *In vitro* cytotoxic activity

Cytotoxic activity of loncastuximab tesirine was assessed *in vitro* as previously described (25). Briefly, cells were exposed to each compound for 96 hours and assayed by MTT [3-(4,5-dimethylthiazolyl-2)-2, 5-diphenyltetrazoliumbromide]. For R-CHOP treatment, cells were exposed for 72 h to 1 μg/mL CHOP + 100 μg/mL rituximab to five different concentrations in serial dilution 1:10. Rituximab was diluted to clinically recommended serum levels (26) and CHOP represented a mix reflecting the clinical ratios of the drugs (27,28) (85%, 4-hydroperoxy-cyclophosphamide; 5.5%, doxorubicin; 0.16%, vincristine; 11.1%, prednisolone). Cells were also exposed in parallel to the PBD dimer SG3199 and of the isotype-control ADC B12-SG3249 (29).

Synergism assessment was done by exposing cells (96 hours) to increasing doses of loncastuximab tesirine and each of the other agents, either alone or in combination, followed by MTT assay. Determination of the Chou-Talalay combination index (CI) was done as previously described (30). Combinations were defined as synergistic (median CTI < 0.9), additive (median CTI, 0-9-1.1) or of no benefit/antagonist (median CTI > 1.1).

### CD19 expression

Absolute cell surface CD19 expression was determined via quantification of the antigen on the surface of lymphoma cell lines using Quantum Simply Cellular (QSC) anti-Human IgG beads (Bangs Laboratories) to create a calibration curve. Antibody Binding Capacity (ABC) values were then normalized to the control isotype antibody B12.

CD19 RNA expression values were extracted from the datasets GSE94669, previously obtained using a targeted RNA-seq approach (HTG EdgeSeq Oncology Biomarker panel), and a microarray-based technology (Illumina HT-12 arrays) (30) and GSE221770, previously produced via total-RNA-Seq (24).

### LyV4.0 CAPP-seq gDNA Assay and variant calling

Genomic DNA was extracted from cell lines using the DNeasy Blood & Tissue Kits (Qiagen, Hilden, Germany). Library preparation started with shearing at least 500 ng of DNA through sonication (Covaris, Woburn, MA) to obtain 100 to 200 bp fragments. The gDNA libraries were then generated with the KAPA Hyper Prep Kit (KAPA Biosystems). The regions of interest (Table S1) were enriched using SeqCap HyperChoice Library probes (NimbleGen; Roche Diagnostics, Jakarta, Indonesia). Libraries were sequenced on the NextSeq500 (Illumina, San Diego, CA) instrument by paired-end sequencing (2 × 150-cycle protocol). A total of 52 multiplexed libraries were simultaneously sequenced in each deep experiment. Sequencing reads in FASTQ format were deduplicated utilizing FastUniq v1.1. The resulting reads were locally aligned to the hg19 human genome assembly using the BWA-MEM v.0.7.17 software with the default settings, and then, sorted, indexed, and assembled into an mpileup file using SAMtools v.1.7. The aligned reads were processed with mpileup using the parameters -A -d 10 000 000. Single nucleotide variations and indels were called in gDNA with the mpileupCNS function of VarScan2 (v.2.2.4) using the parameters min-coverage 1 --min-coverage-normal 1 --min-coverage-tumor 1 --min-var-freq 0 --min-freq-for-hom 0.75 --somatic-p-value 0.05 --min-avg-qual 30 --strand-filter 1 --validation 1 --output VCF. The variant called by VarScan2 were annotated by using the Annovar software (wAannovar https://wannovar.wglab.org/). All the variants affecting coding regions or splice sites were retained in the analysis. All variants were systematically compared to on-line databases to confer the origin of somatic status. Somatic origin of non-synonymous single nucleotide variants (SNV) and/or inframe In/del was confirmed only if was detected as “somatic confirmed” in COSMIC database (https://cancer.sanger.ac.uk/cosmic), without presence in polymorphisms database (https://www.ncbi.nlm.nih.gov/variation/view/ ; https://www.ncbi.nlm.nih.gov/variation/tools/1000genomes/); somatic status was also confirmed by the high damaging prediction score provided by poliphen2 and SIFT online software (http://genetics.bwh.harvard.edu/pph2/ ; https://sift.bii.a-star.edu.sg/www/SIFT4G_vcf_submit.html). For truncating variants, frameshift, splicing variants and/or stop codons, their non-presence in polymorphisms’ database was enough to call variants as somatic. To filter out systematic sequencing errors in gDNA, an in-house database containing all gDNA background allele frequencies across gDNA samples from healthy subjects was used. Based on the assumption that all background allele fractions follow a normal distribution, a Z-test was employed to test whether a given variant in the gDNA differed significantly in its frequency from typical gDNA background at the same position in all gDNA samples, after adjusting for multiple comparisons by Bonferroni test (multiple comparisons corrected p threshold corresponding to alpha of 0.05/[size of the target region in bp × 4 alleles per position]). Variants that did not pass these filters were not further considered. Variants for the resulting candidate mutations were visualized using Integrative Genomics Viewer. Genes mutational levels were correlated with loncastuximab tesirine drug activity quantified as IC_50_ values, by Mann Whitney test with STATA Stata/BE 17.0 (Stata Corporation, College Station, TX). P value for significance was <0.05. For multiple correction analysis, two-stage linear step-up procedure of Benjamini, Krieger and Yekutieli was adopted significance with a threshold of <0.05, using Prism software v8.0 (GraphPad Software La Jolla, CA, USA).

### Immunoblotting

Cells were seeded in T25 flasks at density of 5×10□ per mL and treated for 24 hours with DMSO or single drugs at their 2 times IC_50_ concentrations or with loncastuximab tesirine plus venetoclax, idelalisib or copanlisib. Protein extraction was performed lysing the cells with M-PER (Mammalian Protein Extraction Reagent, ThermoFisher Scientific, Waltham, MA, USA) lysis buffer plus Halt Protease and Phosphatase Inhibitor Cocktail, EDTA-Free (100X) for 30 minutes on ice and then centrifuged at high speed and 4°C for 30 minutes. Protein concentration was determined using the BCA protein assay (Pierce Chemical Co, Dallas, TX, USA) and 30 μg pf total proteins were loaded and separated on a 4-20% gradient SDS-polyacrylamide gel by electrophoresis (SDS-PAGE). Proteins were transferred on nitrocellulose membranes and incubated with primary antibodies overnight, followed by the appropriate horseradish peroxidase–conjugated anti-mouse (NA931V) or anti-rabbit (NA934V) secondary antibodies (GE healthcare, Chicago, IL, USA) for 1 hour at room temperature. Enhanced chemiluminescence detection was done following the manufacturer’s instructions (Amersham Life Science). Luminescence is measured by the Fusion Solo S instrument (Witec AG, Sursee, Switzerland). Finally, protein quantification was performed using the Fusion Solo S instrument (Witec AG). Equal loading of samples was confirmed by probing for vinculin. The antibodies used for the experiment were: anti-Vinculin (Sigma Aldrich cat. n.V9131), anti-AKT (CST-9272), anti-p-AKT (Ser 473) (CST-4060), anti-CD19 (abcam-AB134114), anti-PARP1 (SC-8007), anti-Mcl1 (D35A5) (CST-5453) and anti-Bcl2 (SC-492) as primary antibodies.

### Cell cycle

Cells were seeded in 96 wells-plates at density of 10□ (OCI-LY-3) or 2×10□ (TMD8, VAL, WSU-DLCL2) per wells and subsequently treated with single drugs or with the combination of loncastuximab tesirine plus venetoclax, idelalisib or copanlisib at 2 times IC_50_ concentrations for 96 hours. Cells were fixed with 70% cold ethanol before staining with propidium Iodide (PI) and RNAse treatment. Acquisitions were carried out with a FACSCanto II instrument (BD Biosciences, Allschwil, Switzerland) and data were analyzed using FlowJo software (TreeStar Inc., Ashland, OR, USA).

### Data mining

Statistical analyses were conducted using mainly Prism software v8.0 (GraphPad Software La Jolla, CA, USA). For immunoblotting, cell proliferation, cell death, cell cycle and for apoptotic assay statistical significance was determined by a two-tailed unpaired Student’s t test. A P value < 0.05 was considered statistically significant. The presence of *BCL2* and/or *MYC* translocations and *TP53* inactivation were retrieved from our previous publication (30). Differences in IC_50_ values among lymphoma subtypes were calculated using the Mann-Whitney test. Statistical significance was defined by P values of 0.05 or less.

### In vivo experiments

#### TMD8 xenograft

Mice maintenance and animal experiments were performed under the institutional guidelines established for the Animal Facility at The Institute of Research in Biomedicine (IRB) (license n. TI 49-2018). NOD-SCID mice were obtained from Charles River (Wilmington, MA, USA). Xenografts were established by injecting TMD8 lymphoma cells (15 × 10^6^ cells/mouse, 200 μL of PBS) into the left flanks of female NOD-SCID mice (6 weeks of age, approximately 20 gram of body weight). Treatments started with measurable tumors. Tumor volume (TV) was calculated using the equation V = [length x width x height]/2. During housing and treatments, the animal status was evaluated measuring the Body Condition Score (BSC) (31).

#### JEKO1 xenograft

Eight-week-old female Nod/SCID/IL2-Rg-/-(NSG) mice (Jackson Lab, Bar Harbor, Maine) were injected subcutaneously into both flanks with 10 × 10^6^ Jeko1 cells. Mice were sacrificed according to institutional guidelines (signs of significant disease morbidity such as limb paralysis or greater than 20% weight loss). Animal experiments were approved by and performed in accordance with the guidelines and regulations set forth by the Institutional Animal Care and Use Committee of Case Western Reserve University.

In combination experiments, statistical significances between groups were defined using Mann-Whitney test followed by two-stage step-up (Benjamini, Krieger, and Yekutieli) multiple comparisons, FDR=1%. The coefficient of drug interaction (CDI) (32) was used to assessed the additive (CDI = 1), supra-additive (synergism, CDI < 1) or sub-additive (CDI > 1) effect of the treatment versus the control arms, as previously performed (33). In single treatment experiments, differences between tumor volumes were considered statistically significant using Mann-Whitney test, P<0.05.

## Results

### Loncastuximab tesirine has strong cytotoxic activity in B cell lymphoma cell lines

Loncastuximab tesirine was tested for its anti-proliferative activity across 60 lymphoma cell lines, exposed to the ADC for 96 hours (Table S2). Loncastuximab tesirine had activity in the picomolar range, with a median IC_50_ of 4.1 pM (95% C.I, 2-9.6 pM), among 48 lymphoma cell lines derived from mature B cell lymphomas. Conversely, the anti-proliferative activity of loncastuximab tesirine was over 800-fold lower in nine T cell lymphoma cell lines (median IC_50_ 3.5 nM; 95% C.I, 0.8-11 nM; P<0.0001) (Figure S1). Activity was similar among all the individual B cell lymphoma subtypes with exception of Hodgkin lymphoma models, which were over 600-fold less sensitive to loncastuximab tesirine than the other cell lines (P=0.009) (Table 1). Loncastuximab tesirine exerted its anti-lymphoma activity via induction of apoptosis, as shown in two exemplar cell lines derived from ABC (TMD8) or GCB (VAL) DLBCL (Figure S2).

**Table 1.**
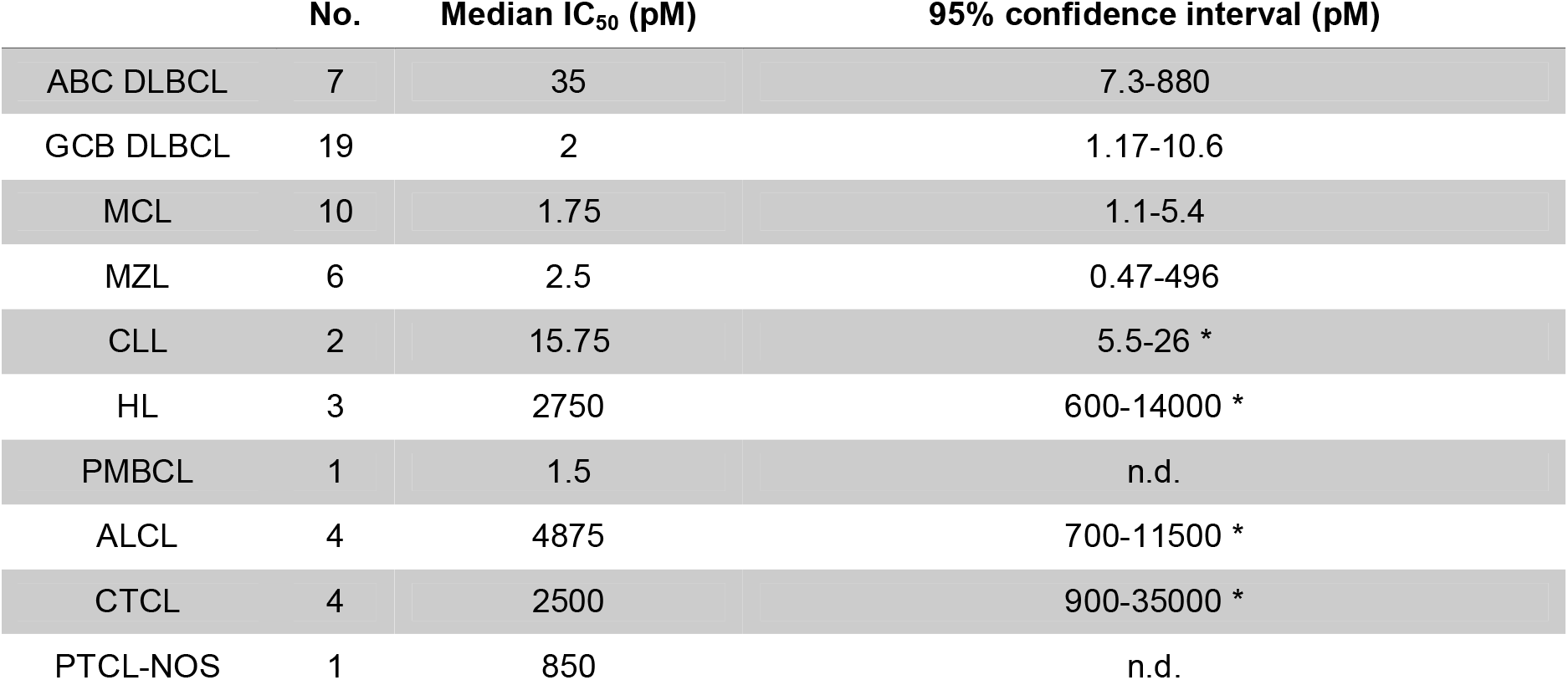
Anti-tumor activity of loncastuximab tesirine in lymphoma cell liness. IC_50_ values obtained after 96 hours treatment. DLBCL, diffuse large B-cell lymphoma; ABC, activated B cell; GCB, germinal center B cell; MCL, mantle cell lymphoma; MZL, marginal zone lymphoma; CLL, chronic lymphocytic leukemia; HL, Hodgkin lymphoma; PMBCL, primary mediastinal large B cell lymphoma; CTCL, cutaneous T cell lymphoma; ALCL, anaplastic large cell lymphoma; PTCL-NOS, peripheral T cell lymphoma-not otherwise specified. n.d., not determined. * Upper confidence limit held at maximum of sample.

The sensitivity to loncastuximab tesirine did not differ between DLBCL cell lines with (n=15) and without (n=11) *BCL2* translocation, or with (n=16) and without (n=7) *TP53* inactivation. Instead, DLBCL cell lines with *MYC* translocation (n=10) versus cell lines without the translocation (n=16) and DLBCL cell lines with (n=7) versus those without (n=19) concomitant *BCL2* and *MYC* translocation (double hit) had lower IC_50_ values (both comparisons, P<0.05) (Figure S3).

The sensitivity to loncastuximab tesirine was also correlated with mutational status obtained from a targeted DNA sequencing designed to cover a variety of different coding genomic regions known to be recurrently mutated in mature B-cell neoplasms (Table S3). After multiple corrections, no somatic mutation was significantly associated to loncastuximab tesirine response.

In parallel, we exposed the cells to an isotype-control ADC (B12-SG3249), which was active in the nanomolar range with no difference between B and T cell lymphoma cell lines: median IC_50_ values were 0.9 nM (95% C.I, 0.7-2.2 nM) and 1.7 nM (95% C.I, 0.8-12 nM), respectively.

Finally, loncastuximab tesirine was tested in three non-human lymphoma cell lines: IC_50_ values were 2 nM and 500 pM in two mouse cell lines, and 175 pM in a canine DLBCL cell line, similar to what was achieved using the isotype-control ADC B12-SG3249, indicating a non-cross species anti-lymphoma activity not driven by CD19 targeting (Table S2).

### CD19 levels correlate with loncastuximab tesirine cytotoxic activity

We then focused on cell lines derived from mature B cell lymphomas to assess whether the levels of CD19 cell surface expression correlated with the anti-tumor activity of loncastuximab tesirine. We measured the absolute CD19 surface expression levels on each cell line (Table S2), and we used additional protein and RNA expression data we had previously obtained on the same panel of cell lines (11,24). We observed that the CD19 expression levels associated with loncastuximab tesirine activity, as demonstrated by the negative correlation between IC_50_ values and CD19 expression values measured both at the cell surface protein level [(absolute quantitation, n=46, r=-0.44, P=0.002; relative quantitation, n=45, r=-0.4, P=0.006]) and RNA level [(microarrays, n=53, r=-0.74, P<0.0001; HTG, n=36, r=-0.5 P=0.002; RNA-Seq, n=44, r=-0.55, P<0.0001] (Figure 1A-E).

**Figure 1.**
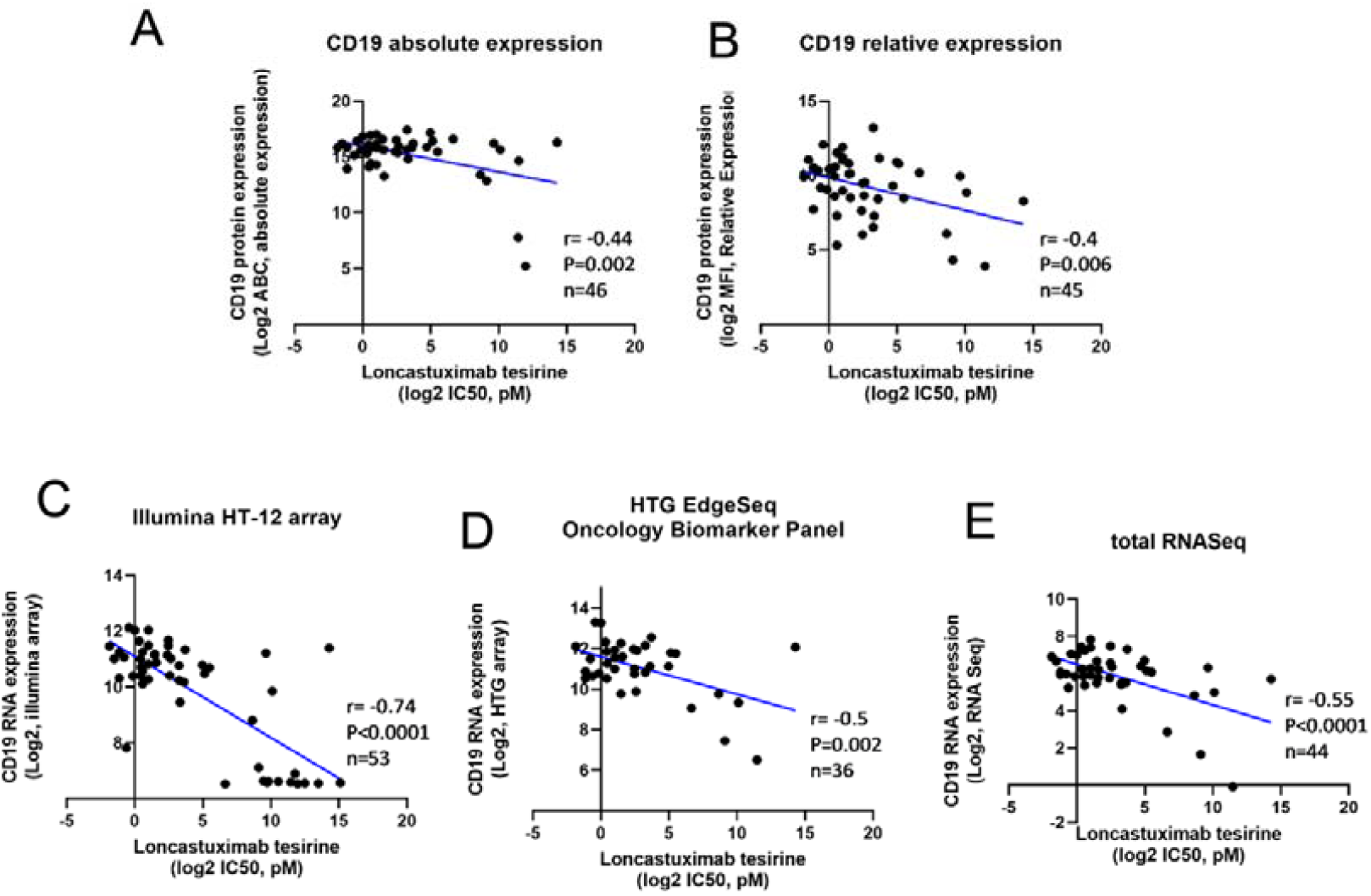
The in vitro anti-proliferative activities of loncastuximab tesirine correlated with CD19 expression. Pearson correlations between loncastuximab tesirine activity with CD19 protein absolute and relative expression, CD19 transcript measured with the Illumina HT-12 arrays, HTG biomarker panel and total RNASeq data.

Based on the above-mentioned association between the presence of *MYC* translocation and loncastuximab tesirine lower IC_50_ values (i.e., higher sensitivity), we explored the possible relationships between CD19 and MYC expression levels in DLBCL cells. Nor CD19 surface protein expression levels nor CD19 RNA levels differed between cell lines with or without MYC translocation (single genetic event or together with BCL2 translocation) (Figure S4A-C). Similarly, CD19 and MYC levels were not correlated (Figure S4D-E). Finally, MYC RNA levels were negatively correlated with loncastuximab tesirine IC_50_ values (R=-0.35), but without reaching a statistical significance (P=0.089) (Figure S4F).

### The cytotoxic activities of loncastuximab tesirine’s warhead SG3199 is not affected by lymphoma subtype but differs based on the presence of genetic lesions

All cell lines were exposed to loncastuximab tesirine’s warhead SG3199 (Table S2). The median IC_50_ value was 0.85 pM (95% C.I, 0.69-1.14) across all 60 lymphoma cell lines. Differently from what was observed with loncastuximab tesirine, the activity of SG3199 did not differ between B and T cell lymphomas (Table 2, Figure S5), and there was no correlation between the SG3199 IC_50_ values and CD19 expression values (Figure S6). SG3199 was more potent than the ADC. The difference in terms of IC_50_ values between SG3199 and loncastuximab tesirine was statistically significant both considering all cell lines (P<0.0001) or cell lines derived from B cell lymphomas (P<0.0001) (Figure S7).

**Table 2.** Anti-tumor activity of the SG3199 warhead in lymphoma cell lines. IC_50_ values obtained after 96 hours treatment. DLBCL, diffuse large B-cell lymphoma; ABC, activated B cell; GCB, germinal center B cell; MCL, mantle cell lymphoma; MZL, marginal zone lymphoma; CLL, chronic lymphocytic leukemia; HL, Hodgkin lymphoma; PMBCL, primary mediastinal large B-cell lymphoma; CTCL, cutaneous T cell lymphoma; ALCL, anaplastic large cell lymphoma; PTCL-NOS, peripheral T cell lymphoma-not otherwise specified. n.d., not determined. * Upper confidence limit held at maximum of sample.

|  | No. | Median IC<sub>50</sub> (pM) | 95% confidence interval (pM) |
| --- | --- | --- | --- |
| ABC DLBCL | 7 | 1.17 | 0.63-7.85 |
| GCB DLBCL | 19 | 1.14 | 0.75-1.53 |
| MCL | 10 | 0.53 | 0.53-1.66 |
| MZL | 6 | 0.53 | 0.53-0.85 |
| CLL | 2 | 0.83 | 0.53-1.14* |
| HL | 3 | 4.97 | 0.85-29.24* |
| PMBCL | 1 | 0.56 | n.d. |
| ALCL | 4 | 2.34 | 0.85-17.54* |
| CTCL | 4 | 1.59 | 0.53-23.39* |
| PTCL-NOS | 1 | 0.53 | n.d. |

The sensitivity to SG3199 appeared reduced in DLBCL cell lines with *TP53* inactivation when compared to *TP53* wt cell lines (P<0.001) (Figure S8A). The presence of the *BCL2* translocation *per se* did not affect the sensitivity to SG3199 (Figure S8B). Like what was observed with loncastuximab tesirine, SG3199 was more active in DLBCL bearing *MYC* translocation as single event or concomitant with *BCL2* translocation (P<0.05) (Figure S8C-D). No correlation was observed between sensitivity to SG3199 and MYC RNA levels (Figure S9).

### The cytotoxic activities of loncastuximab tesirine and its warhead SG3199 are highly correlated

The cytotoxic activity of loncastuximab tesirine and its warhead were highly positively correlated among all the cell lines (r=0.60, P<0.0001) and within the cell lines derived from mature B cell lymphomas (r=0.63, P<0.0001) (Figure 2). Most of the cell lines that were less sensitive to the ADC (IC_50_ values higher than the 75^th^ percentile, i.e. 768 pM), but sensitive to SG3199 (IC_50_ values lower than the 75^th^ percentile, i.e. 2.9 pM) were represented by the CD19 negative models (T cell lymphomas, HL) and the non-human lymphomas. Some cell lines, such as the splenic MZL VL51, were highly sensitive to the warhead, but, due to low CD19 expression, high loncastuximab tesirine IC_50_ (VL51 IC_50_ >100 fold the median B cell lymphoma cell lines IC_50_). There were a few cell lines, especially the mantle cell lymphoma (MCL) REC1 and the DLBCL Pfeiffer and U2932, which had IC_50_ values higher than the 75^th^ percentile for both loncastuximab tesirine and SG3199 warhead, suggestive of a primary resistance to the warhead. The GCB DLBCL cell line SU-DHL-6 was sensitive to loncastuximab tesirine but resistant to its warhead SG3199. We confirmed that the anti-tumor activity was driven by the activity of the antibody itself rather than by the antibody complexed to the toxin (Figure S10).

**Figure 2.**
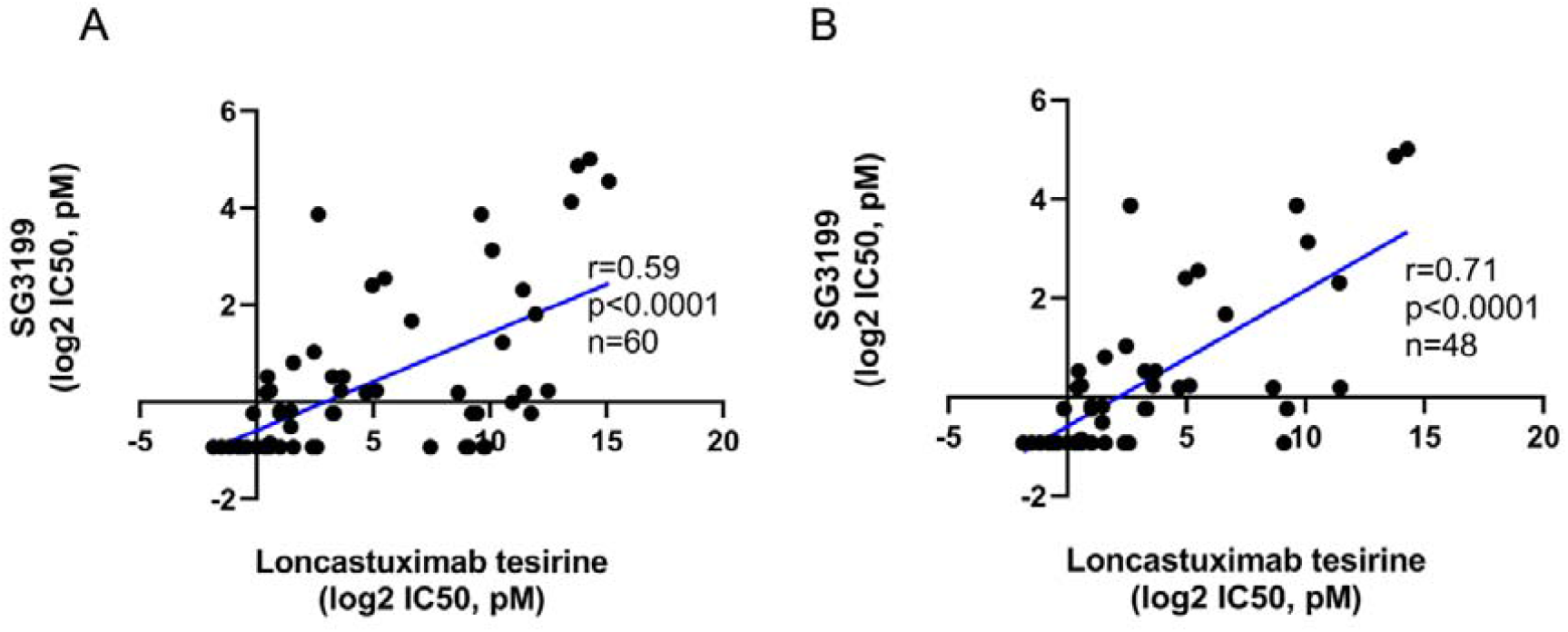
Correlation between the activity of loncastuximab tesirine and its warhead SG3199. Pearson correlations between loncastuximab tesirine and SG3199 IC_50_ values across all cell lines (A) and in cell lines derived from B cell lymphomas and HL (B).

### The cytotoxic activity of loncastuximab tesirine is correlated with the cytotoxicity of other CD19 targeting ADCs

We exploited data previously produced in our laboratory on the same panel of cell lines with two CD19 targeting ADCs, coltuximab ravtansine (SAR3419), comprising the maytansinoid microtubule disruptor DM4, and huB4-DGN462, incorporating a DNA-alkylating payload (11). The pattern of activity of loncastuximab tesirine correlated with both coltuximab ravtansine (r=0.38, P=0.01) and huB4-DGN462 (r=0.6, P<0.0001) (Figure S11). Loncastuximab tesirine was more potent than both huB4-DGN462 (P=0.034) and coltuximab ravtansine (P<0.0001), although the exposure time previously used for the two additional ADCs was shorter (72 vs 96 hours). REC1, Pfeiffer and U2932, the cell lines most resistant to loncastuximab tesirine, were also resistant to huB4-DGN462 and coltuximab ravtansine.

### The pattern of cytotoxic activity of loncastuximab tesirine and R-CHOP are correlated

We exposed DLBCL cell lines (n=26) to the *in vitro* equivalent of R-CHOP (rituximab, cyclophosphamide, doxorubicin, vincristine, prednisone) (Table S3). The IC_50_ values obtained with R-CHOP correlated with the IC_50_ values of both loncastuximab tesirine (r=0.667, P<0.001) and SG3199 (r=0.421, P=0.032) (Figure 3). Some cell lines presented a reduced sensitivity to R-CHOP (IC_50_ values higher than the 75^th^ percentile, i.e., 0.9 μg/mL), but were very sensitive to loncastuximab tesirine and to its warhead. A few cell lines (Pfeiffer, U2932, SU-DHL-16, SU-DHL-2) were less sensitive to both loncastuximab tesirine and R-CHOP.

**Figure 3.**
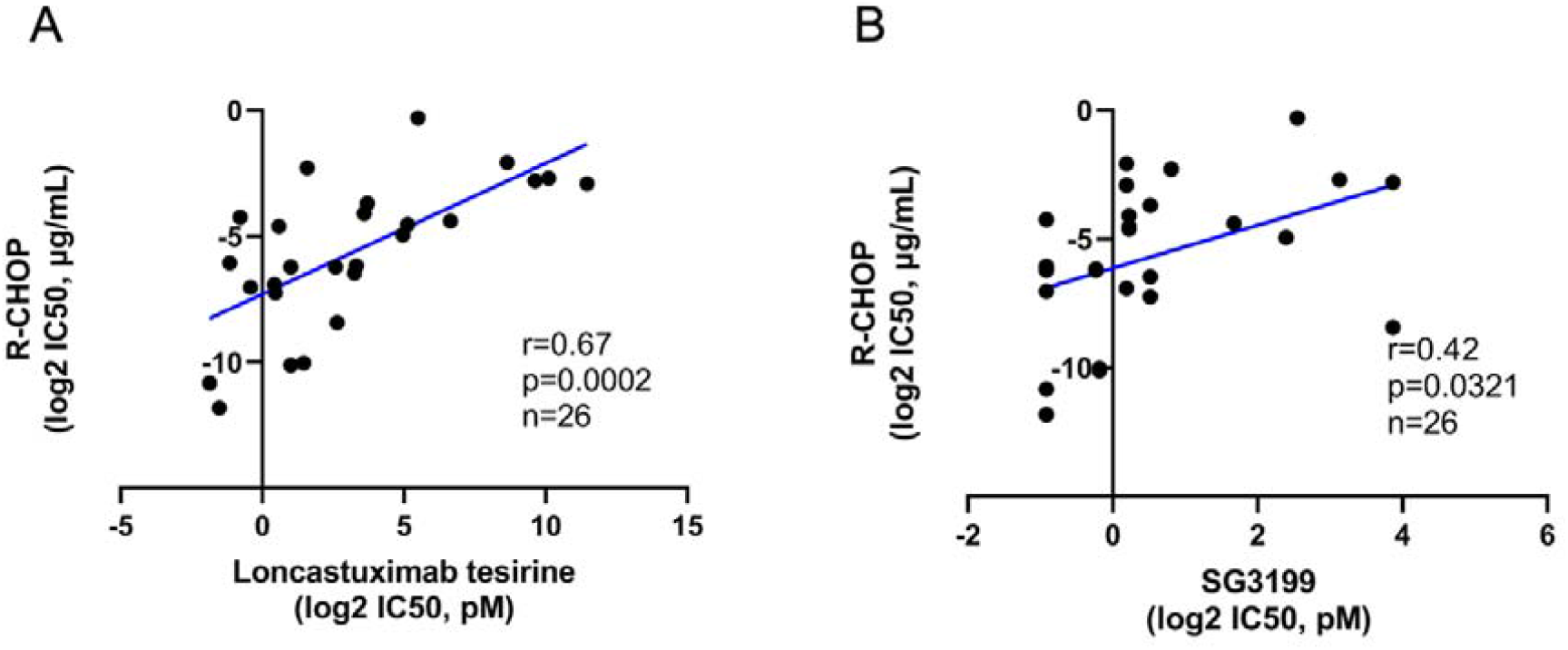
Correlation between the activity of loncastuximab tesirine or its warhead SG3199 and R-CHOP in DLBCL cell lines. Pearson correlations between R-CHOP and loncastuximab tesirine (A) or SG3199 (B).

### Loncastuximab tesirine-based combinations appear beneficial *in vitro*

We explored the potential benefit of combining loncastuximab tesirine with drugs having an established role in the treatment of lymphoma patients. We tested these combinations in two GCB DLBCL cell lines (VAL and WSU-DLCL2), two ABC DLBCL (TMD8 and OCI-LY-3) and two MCL (JEKO1 and JVM2). The combination partners were the BCL2 inhibitor venetoclax, the PI3Kδ inhibitor idelalisib, the PIK3α/δ inhibitor copanlisib, the anti-CD20 monoclonal antibody rituximab, the chemotherapy agent bendamustine, and the PARP inhibitor olaparib (Table 3). The combinations of loncastuximab tesirine with the proteasome inhibitor bortezomib, the BTK inhibitor ibrutinib, and the immunomodulator lenalidomide, were only tested the ABC DLBCL cell lines as these drugs are only used for the treatment of ABC DLBCL.

**Table 3.** Loncastuximab tesirine containing combinations in DLBCL and MCL cell lines; ABC DLBCL, activated B cell like diffuse large B cell lymphoma; GCB DLBCL, germinal center B cell like diffuse large B cell lymphoma; MCL, mantle cell lymphoma.

| Combination Partner | Histology | Cell line | Median Chou-Talalay combination index | 95% confidence interval |
| --- | --- | --- | --- | --- |
| Venetoclax | ABC DLBCL | OCI-Ly-3 | 0.48 | 0.3-0.6 |
|  | ABC DLBCL | TMD8 | 0.63 | 0.48-1.17 |
|  | GCB DLBCL | VAL | 0.75 | 0.66-0.91 |
|  | GCB DLBCL | WSU-DLCL2 | 0.4 | 0.19-1.86 |
|  | MCL | JVM2 | 0.37 | 0.25-0.69 |
|  | MCL | JEKO-1 | 0.88 | 0.37-1.01 |
| Copanlisib | ABC DLBCL | OCI-Ly-3 | 0.53 | 0.41-0.68 |
|  | ABC DLBCL | TMD8 | 1.07 | 0.52-1.22 |
|  | GCB DLBCL | VAL | 0.84 | 0.63-0.93 |
|  | GCB DLBCL | WSU-DLCL2 | 1.56 | 0.87-1.80 |
|  | MCL | JVM2 | 0.19 | 0.09-0.47 |
|  | MCL | JEKO-1 | 0.75 | 0.41-1.05 |
| Idelalisib | ABC DLBCL | OCI-Ly-3 | 0.1 | 0.07-0.22 |
|  | ABC DLBCL | TMD8 | 0.9 | 0.41-1.24 |
|  | GCB DLBCL | VAL | 0.86 | 0.67-1.22 |
|  | GCB DLBCL | WSU-DLCL2 | 0.5 | 0.30-0.75 |
|  | MCL | JVM2 | 0.42 | 0.36-0.79 |
|  | MCL | JEKO-1 | 1.32 | 0.68-1.65 |
| Bendamustine | ABC DLBCL | OCI-Ly-3 | 1 | 0.7-1.75 |
|  | ABC DLBCL | TMD8 | 0.6 | 0.35-1.92 |
|  | GCB DLBCL | VAL | 0.89 | 0.59-1.14 |
|  | GCB DLBCL | WSU-DLCL2 | 0.62 | 0.51-0.83 |
| Bortezomib | ABC DLBCL | OCI-Ly-3 | >3 | >3 |
|  | ABC DLBCL | TMD8 | 1.13 | 0.86-1.57 |
| Ibrutinib | ABC DLBCL | OCI-Ly-3 | 0.76 | 0.39-0.9 |
|  | ABC DLBCL | TMD8 | 1.07 | 0.96-1.25 |
| Lenalidomide | ABC DLBCL | OCI-Ly-3 | 1.58 | 0.82-4.08 |
|  | ABC DLBCL | TMD8 | 1.88 | 0.93-2.36 |
| Olaparib | ABC DLBCL | OCI-Ly-3 | 1.41 | 0.99-2.06 |
|  | ABC DLBCL | TMD8 | 0.95 | 0.41-1.29 |
|  | GCB DLBCL | VAL | 0.86 | 0.67-1.22 |
|  | GCB DLBCL | WSU-DLCL2 | 0.5 | 0.30-0.75 |
| Rituximab | ABC DLBCL | OCI-Ly-3 | >3 | >3 |
|  | ABC DLBCL | TMD8 | >3 | >3 |
|  | GCB DLBCL | VAL | 0.09 | 0.06-0.14 |
|  | GCB DLBCL | WSU-DLCL2 | >3 | >3 |

In DLBCL, the most beneficial combinations were loncastuximab tesirine plus venetoclax or idelalisib, with synergism achieved in all the models tested, followed by the ADC plus bendamustine, copanlisib or olaparib. Synergism was also observed in one (OCI-Ly-3) of the two ABC DLBCL cell lines tested with loncastuximab tesirine plus ibrutinib. Combination with rituximab was synergistic in one cell line (VAL). No advantage was seen in combining loncastuximab tesirine with bortezomib or lenalidomide. In MCL the most beneficial combinations were observed with venetoclax and copanlisib, with synergism in two out of two cell lines. The addition of idelalisib was synergistic in only one cell line (JVM2).

The effect on cell cycle was investigated in four DLBCL cell lines (TMD8, OCI-LY-3, VAL and WSU-DLCL2) treated with loncastuximab tesirine and the most promising targeted agents, i.e., venetoclax, idelalisib and copanlisib, as single agents and in combination, after 96 hours of treatments. In all the DLBCL cell lines the increase in sub-G0, compatible with cell death, was higher compared to the control in loncastuximab tesirine treated cells either as single agent or in combination (Figure S12). Treatments with venetoclax, idelalisib and copanlisib as single agents also induced an increase in sub-G0 in WSU-DLCL2 and VAL and OCI-LY-3. In TMD8 an increase in sub-G0 was only observed in single copanlisib treatment.

To understand the mechanism sustaining the benefit given by loncastuximab tesirine when combined with venetoclax, idelalisib or copanlisib, specific signaling pathways were explored by immunoblotting in TMD8 (ABC DLBCL) and WSU-DLCL2 (GCB DLBCL) cell lines (Figure 4, Figure S13). CD19 downregulation was observed in both cell lines following treatment with loncastuximab tesirine alone after 24 hours of treatment, and the downregulation increased further when loncastuximab tesirine was combined with each of the three drugs. The levels of pAKT were reduced in cells treated with the PI3K inhibitors idelalisib and copanlisib as single agents or in combination with loncastuximab tesirine (Figure 4, Figure S7). The levels of the anti-apoptotic protein MCL1 were downregulated by treatment with loncastuximab tesirine in the TMD8 cell line, and the effect was maintained in the combinations. In the WSU-DLCL2 cell line, MCL1 levels were also down-regulated in cells treated with loncastuximab tesirine and with the PI3K inhibitors as single agents and in combination. In both cell lines, exposure to venetoclax upregulated MCL1, which was reduced in combination with loncastuximab tesirine. Cleaved PARP1, a marker of apoptosis, was slightly increased when loncastuximab tesirine was combined with the other three drugs.

**Figure 4.**
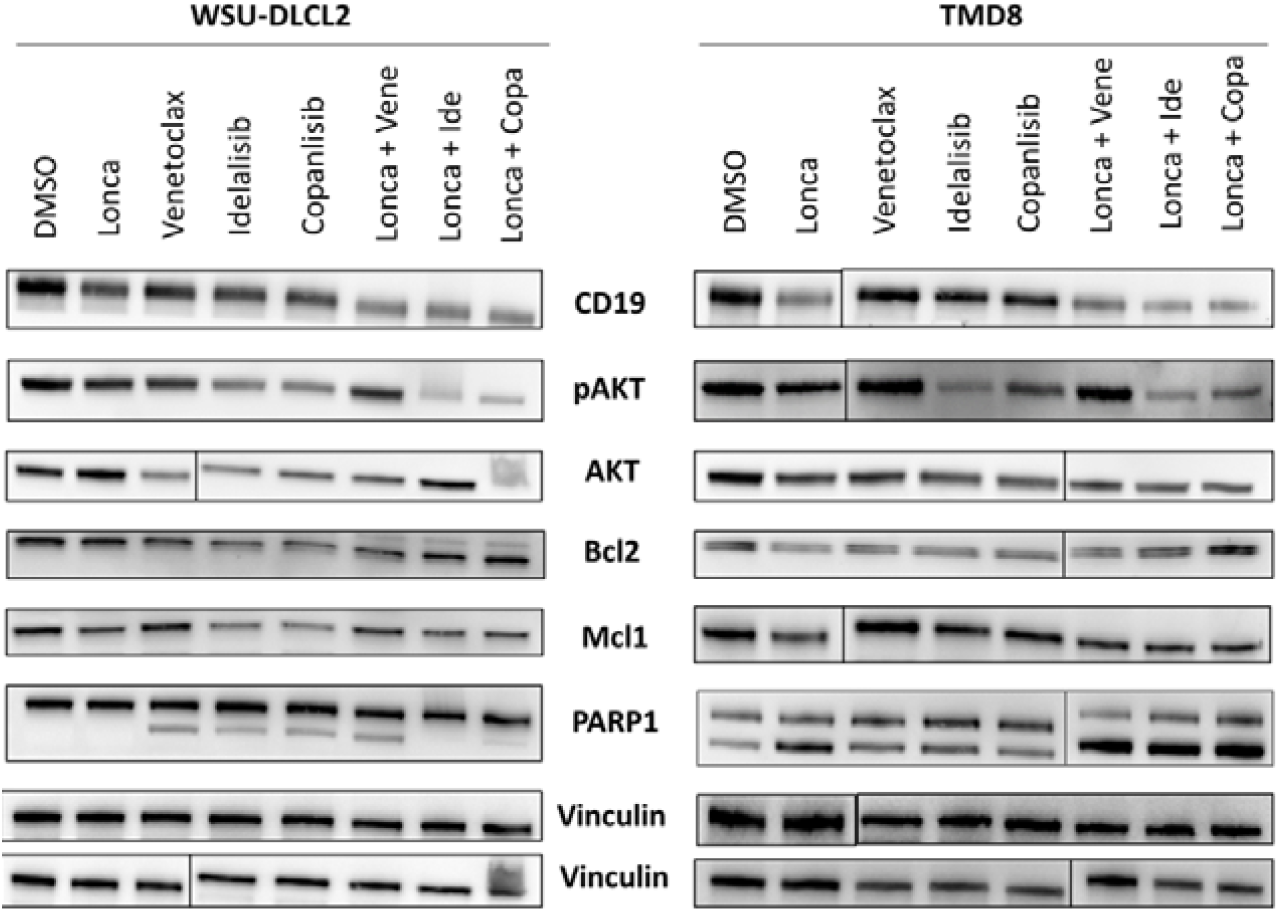
Proteins modulated after exposure to loncastuximab tesirine. Representative immunoblots from one ABC DLBCL (TMD8) and one GCB DLBCL (WSU-DLCL2) cell lines treated for 24 hours with drugs as single or the combination of loncastuximab tesirine with venetoclax, idelalisib and copanlisib at concentrations corresponding to 2 times the IC_50_ values.

### Loncastuximab tesirine-based combinations are beneficial *in vivo*

An ABC DLBCL xenograft (TMD8 cell line) was used to validate the combination of loncastuximab tesirine with copanlisib in vivo. We first evaluated both compounds as single agents to define the doses to be combined. Mice (n.= 5 per group) bearing subcutaneously (sc) TMD8 xenografts were treated with control (PBS, iv), three different doses of loncastuximab tesirine (0.1 mg/Kg vs 0.3 mg/Kg vs 0.6 mg/Kg; iv qd x 1), two different doses of the non-binding control ADC B12-SG3249 (0.3 mg/Kg vs 0.6 mg/Kg; iv qd x 1) (Figure S14), or two different schedules of copanlisib (13 mg/Kg, iv; 1 day on/6 days off vs 2 days on/5 days off) (Figure S15). Based on the observed dose-dependent anti-tumor activity for both loncastuximab tesirine and copanlisib, we defined the sub-active schedules of the drugs to be used in the combination study. Thus, the combination experiment included animals treated with a single dose of loncastuximab tesirine (0.3 mg/Kg iv; day 1; n.=7), or copanlisib (13 mg/kg iv, 1 day on/6 days off; day 1, 8; n.=7) as single agents or in combination (n.=9) (Figure 5A). As control, a group of mice was treated with vehicle (PBS) or with non-binding control ADC B12-SG3249 at 0.3 mg/kg (n.=4 each, iv, qd x 1; day 1). The combination of loncastuximab tesirine with the PIK3α/δ inhibitor copanlisib decreased tumor volume compared to vehicle, isotype control and single treatments (Area Under the Curve, AUC, combination=1.115; vehicle=3.702; B12-SG3249=2.883; copanlisib=1,952; loncastuximab tesirine=2,032). After day 1, the anti-tumor effect of the combination was always superior to the copanlisib (q<0.001), loncastuximab tesirine (q<0.001), vehicle (q<0.001), and ADC isotype control (q<0.001) treatment. The anti-tumor activity of B12-SG3249 did not differ from vehicle alone (q>0.1) or versus the other single agent treatments. In terms of tumor weight, the effect of the combination was superior to the single agents (p=0.003; combo vs copanlisib, p<0.0001; combo vs loncastuximab tesirine, p<0.0001; combo vs ADC isotype, p=0.003) (Figure S10A). The combination presented an additive/slight synergistic CDI at day 38 (CDI=0.959 using B12-SG3249 as control; CDI=1.03 using merged vehicle and B12GS3249 as control). No toxicities were observed with single agents or combinations in this setting.

**Figure 5.**
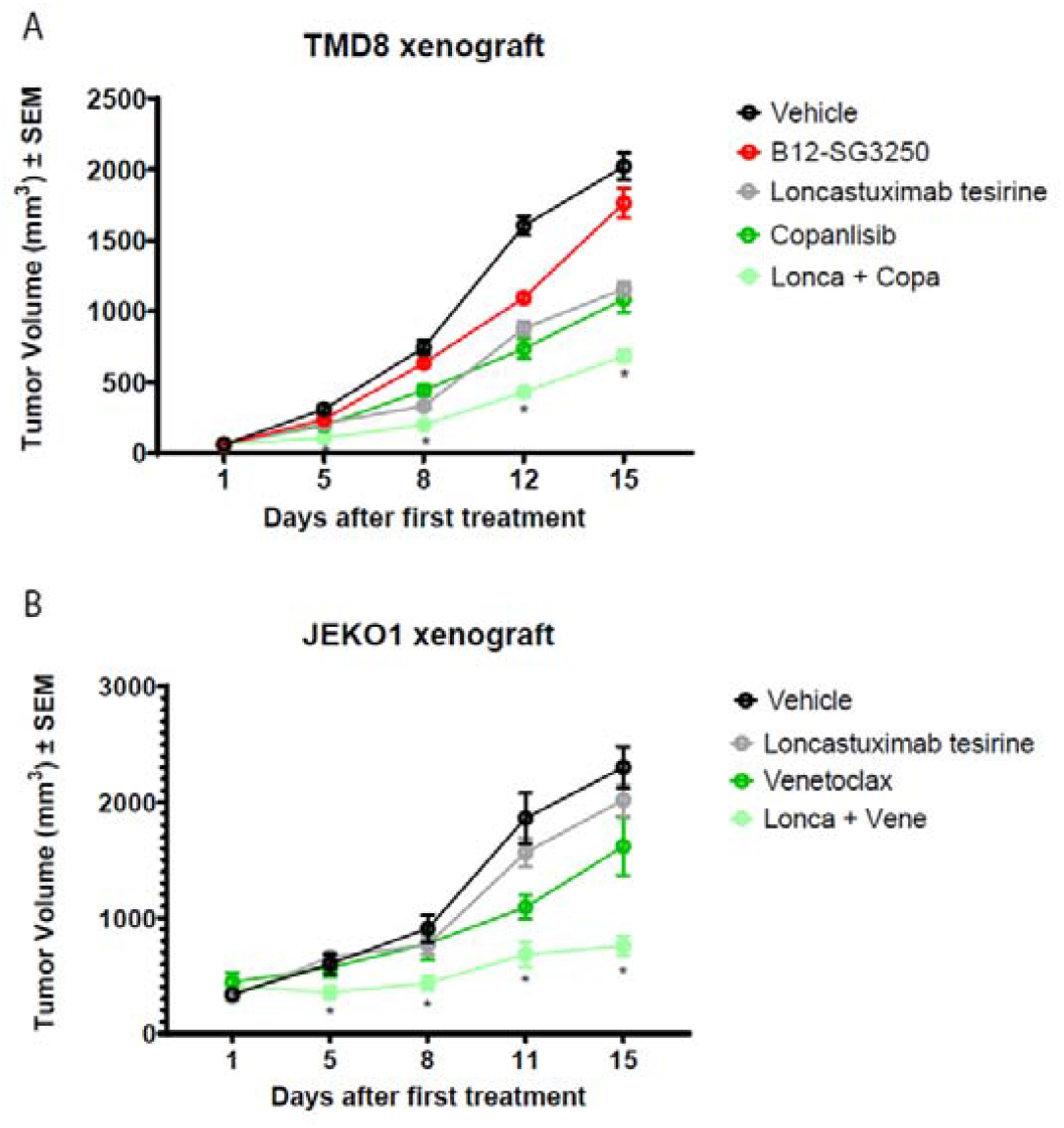
The combination of loncastuximab tesirine plus copanlisib and venetoclax is *in vivo* superior to single agents in ABC DLBCL and MCL xenograft model. (A) NOD-SCID Mice were sc injected with TMD8 and treated (n.=9 per group) with loncastuximab tesirine and of copanlisib as single agents and in combination, and, as control with vehicle (PBS) or the non-binding control ADC B12-SG3249 (n.=4 per group). * q values < 0.01 of combo vs all other groups (vehicle, B12-SG3249, loncastuximab tesirine, copanlisib) was determined by Mann-Whitney test followed by two-stage step-up (Benjamini, Krieger, and Yekutieli) multiple comparisons, FDR(q)=0.01. (B) NSG mice were sc injected with JEKO1 and treated (n.=8 per group) with loncastuximab tesirine and of venetoclax as single agents and in combination, and PBS as control. Y-axis average tumor volume expressed in mm^3^ ± SEM. * q values < 0.01 of combo vs vehicle and loncastuximab tesirine group, as determined by Mann-Whitney test followed by two-stage step-up (Benjamini, Krieger, and Yekutieli) multiple comparisons, FDR(q)=0.01.

A MCL xenograft model (JeKo1) was used to assess the combination of loncastuximab tesirine with venetoclax. Mice (n=4 per group) were treated with a single injection of loncastuximab tesirine (1 mg/kg iv), and/or venetoclax (100mg/kg PO daily 5 days a week), or vehicle at day 16 after cells injection when all tumors were palpable. Tumor volume was decreased in the combination arm compared to vehicle and single treatments (AUC, combination=2,059; vehicle=4,692; venetoclax=3,470; loncastuximab tesirine=4,162), and the anti-tumor effect was statistically significant compared to the vehicle and loncastuximab tesirine (q<0.001) as single agent at all time points but day1 (Figure 5B, Figure S10B). A synergistic coefficient of drug interaction (CDI=0.53) was calculated for this combination at the end of the experiment (day 15).

## Discussion

Here, we shown that i) the CD19 targeting ADC loncastuximab tesirine has a strong cytotoxic activity in a large panel of cell lines derived from B cell lymphomas, ii) its *in vitro* activity correlated with CD19 expression level, and that iii) there is benefit of adding loncastuximab tesirine to other agents, especially BCL2 and PI3K inhibitors. We also showed the similarities and differences in terms of activity with its warhead, other CD19-targeting ADCs and with R-CHOP.

These findings extend the initial preclinical data (10), confirming loncastuximab tesirine cytotoxic activity in mature B cell lymphomas. In the initial publication, only a weak trend was observed between anti-tumor activity of loncastuximab tesirine and CD19 expression levels across ten cell lines that also included CD19-negative cells (10). Here, we expanded the number of cell lines analyzed and even when focusing on B cell lymphoma models only, we observed a significant correlation between the activity of loncastuximab tesirine and CD19 expression on cell surface as well as CD19 RNA levels, using multiple platforms including one specifically designed for the analysis of formalin-fixed paraffin embedded clinical specimens (34). So far, immunohistochemistry (IHC) performed on tumor samples from the loncastuximab tesirine phase 1 and the phase 2 trials have not demonstrated a correlation between CD19 expression and ORR, with patients with extremely low or no detectable CD19 IHC expression responding to loncastuximab tesirine (20,35). However, our data in cell lines and the observation that measuring the CD19 surface density in addition to the IHC expression improves the response prediction (35) suggest that more sensitive measurements of CD19 in clinical specimens might be useful to predict the type and the duration of response of patients treated with loncastuximab tesirine.

Besides loncastuximab tesirine, we also tested its warhead, SG3199, on all the cell lines. As expected, SG3199 did not show any correlation with CD19 expression, and it was equally active in CD19-positive and CD19-negative cell lines. Interestingly, the anti-tumor activity of loncastuximab tesirine correlated with the intrinsic sensitivity of the cell lines to SG3199. Indeed, we could identify three different groups of cell lines. One group of cell lines that were highly sensitive to both loncastuximab tesirine and SG3199 and these presented the highest CD19 expression. A second group of cell lines sensitive to the warhead but not to the ADC (IC_50_ values higher than the 75^th^ percentile). These included the models not derived from human B cell lymphomas, but also B cell lymphomas with low CD19 expression. One example of these was the VL51 cell line, derived from splenic MZL. Interestingly, we recently reported that VL51 derivatives with resistance to PI3K and BTK inhibitors, acquired after months of exposure to idelalisib or ibrutinib, present higher CD19 expression levels than the parental cells and an increased sensitivity to loncastuximab tesirine (36) and anti-CD19 chimeric antigen receptor (CAR) T cells (37), further indicating the importance of CD19 expression levels. A third group of cell lines was characterized by IC_50_ values of both SG3199 and loncastuximab tesirine higher than the 75^th^ percentiles, indicative of a low sensitivity to the agents and of an intrinsic resistance to the PBD warhead.

There was no effect of histology, DLBCL cell of origin, *TP53* or *BCL2* genes status on the *in vitro* cytotoxic activity of loncastuximab tesirine. Among DLBCL cell line models, we discovered an association between *MYC* translocation, as single event or together with BCL2 translocation, and a higher sensitivity to loncastuximab tesirine and to its warhead, which might be sustained by the interplay between MYC-induced replication stress and the SG3199-induced DNA interstrand cross-links (38-40). The clinical relevance of this finding remains unclear. Interestingly, in the phase 2 study, cases with *MYC* translocation were as sensitive as the remaining patients, suggesting that even this group of otherwise poor outcome patients can indeed benefit from the ADC (41).

The cytotoxic activity of PBD dimers can occur via TP53-independent and TP53-dependent mechanisms (39), and we observed a decreased activity of SG3199 in *TP53*-inactive DLBCL cell lines when compared with the wt *TP53* models. Although this difference was not observed when cells were exposed to loncastuximab tesirine, it suggests that payloads with an alternative mechanism of action might work better in the context of an inactive TP53.

Next, we compared the activity of loncastuximab tesirine against all cell lines with the activity of R-CHOP, which is used in the first-line treatment of DLBCL. Loncastuximab tesirine was clearly more active in many cell lines with low/moderate sensitivity to R-CHOP, but the anti-tumor activity of loncastuximab tesirine and of its warhead showed significant correlation with the activity of R-CHOP. Indeed, there were cell lines very sensitive to all treatments and cell lines showing resistance to R-CHOP as well as to loncastuximab tesirine and its warhead. We took advantage of a previous study (11), and we compared the activity of loncastuximab tesirine to coltuximab ravtansine and huB4-DGN462, two other CD19 targeting ADCs which were analyzed using the same panel of cell lines. Interestingly, despite a higher potency, the cytotoxic activity of loncastuximab tesirine correlated with both coltuximab ravtansine and huB4-DGN462. The correlation was higher with the latter ADC, which is *in vitro* and *in vivo* more potent than coltuximab ravtansine (11), and carries the DNA-alkylating agent indolinobenzodiazepine pseudodimer DGN462 as warhead (11), rather than the maytansinoid microtubule disruptor N2′-deacetyl-N2′-(4-mercapto-4-methyl-1-oxopentyl) (DM4 or ravtansine) (42), present in coltuximab ravtansine. This observation and the comparison with R-CHOP highlight the importance of finding novel treatment modalities, including new active molecules as payloads.

We combined loncastuximab tesirine with other anti-lymphoma agents to identify potentially active combinations that may provide better outcomes for patients. In DLBCL, the loncastuximab tesirine – based combinations that were synergistic in most cell lines included those with the BCL2 inhibitor venetoclax, PI3K inhibitors (idelalisib, copanlisib) and with chemotherapy agent bendamustine, followed at less extent by the PARP inhibitor olaparib, the BTK inhibitor ibrutinib and the anti-CD20 monoclonal antibody rituximab. The *in vitro* findings with venetoclax and the PI3K inhibitors were extended to MCL cell lines and additionally confirmed in vivo. While venetoclax has been extensively combined with small molecules, much less data are available regarding the combination with ADCs. Synergy with venetoclax has been previously reported for two ADCs bearing microtubule targeting agents as payloads, the CD79B targeting polatuzumab vedotin and the CD205 targeting MEN1309. Exposure to both agents, containing MMAE and DM4, respectively, caused down-regulation of MCL1 (43,44), due to protein degradation via the ubiquitin/proteasome system (43). Also alkylating agents have been shown to induce proteasome-mediated degradation of MCL1 (45) hence we anticipate a similar mechanism of action mediated by loncastuximab tesirine via its SG3199 payload and sustaining the observed synergism with venetoclax. The combination of loncastuximab tesirine and venetoclax is currently being explored in a phase 1 study (NCT05053659). No trial is currently exploring the combination of loncastuximab tesirine with PI3K inhibitors, which based on the *in vitro* and *in vivo* anti-tumor activity, appears promising. The novel highly specific PI3Kδ inhibitors seem to have an improved toxicity profile (46,47), which might overcome the problems observed with first generation compounds (48).

The combination of loncastuximab tesirine with ibrutinib, also supported by other preclinical work (49), has been clinically evaluated with results reported in R/R DLBCL or MCL (50). The toxicity was manageable, and the overall response rates were 67% in non-GCB DLBCL, 20% in GCB DLBCL, and 86% in MCL (50).

The benefit of combining loncastuximab tesirine with a PARP inhibitor could lead to novel clinical opportunities. The observed benefit of combining a PBD-based ADC with PARP inhibitor is in line with data reported especially in BRCA-deficient solid tumor models (51-53). Interestingly, the GCB DLBCL marker LMO2 inhibits BRCA1 recruitment to DNA double-strand break in DLBCL cells, causing a BRCA1-deficiency-like phenotype and sensitizing DLBCL cells to PARP inhibition (54). Indeed, we observed synergism in the GCB DLBCL cells but only an additive effect in one of the two ABC DLBCL models. PARP inhibitors have been explored in lymphoma patients (55). In particular, the PARP inhibitor veliparib has shown evidence of clinical activity, including complete remissions, and safety in combination with bendamustine plus or minus rituximab (55,56).

Since there are multiple CD19 targeting therapeutic modalities available that share CD19 loss as one of the mechanisms of resistance (57), it will be crucial to define the best sequencing or prioritization strategy for the use of these agents (58-61). In this context, our data clearly support the further development of loncastuximab tesirine as single agent and in combination for patients affected by mature B cell neoplasms. The results also highlight the importance of CD19 expression, and the existence of lymphoma populations characterized by resistance to multiple therapies.

## Acknowledgements

This project was partially supported by research funds from ADC Therapeutics, the Swiss National Science Foundation grant SNSF 310030_197466, the Swiss Cancer Research grant KFS-4713-02-2019. NM was supported by a Ph.D. Fellowship of the NCCR RNA & Disease, a National Centre of Competence in Research funded by the Swiss National Science Foundation (grant numbers 182880, 205601). We thank our Colleagues Dr Antonella Zucchetto and Dr Valter Gattei (Aviano, Italy) for helpful discussion.

## Author Contributions

CT: performed experiments, performed data mining, interpreted data, co-wrote the manuscript.

FS, EG: performed experiments, interpreted data.

LC: performed data mining.

DW, EC, AJA, GG, LS: performed experiments. NM, AB: performed experiments and data mining.

AB, DR: performed targeted DNA sequencing and data mining. EZ, AS: provided advice.

PHVB, FZ: co-designed the study, provided reagents, supervised the study.

FB: co-designed the study, performed data mining, interpreted data, supervised the study and co-wrote the manuscript.

All authors reviewed and accepted final version of the manuscript.

