## Supplementary Figures and Tables for "Targeting CD19-positive lymphomas with the antibody-drug conjugate (ADC) loncastuximab tesirine: preclinical evidence as single agent and as combinatorial approach"

### Supplementary Tables

**Table S1**. **Regions specifically sequenced by the probes in the targeted DNA-sequencing data.** Each row delineates the primary target of genomic spaces covered by the probes according to Hg19 genome reference.

**Table S2. $IC_{50}$ values obtained in lymphoma cell lines after 96 hours exposure to loncastuximab tesirine, SG3199 and isotype-control ADC (B12-C220-SG3249), in addition to CD19 surface protein expression in lymphoma cell lines as measured following an absolute fluorescence quantitation with Quantum Simply Cellular microspheres using rB4v1.2 antibody.** DLBCL, diffuse large B-cell lymphoma; ABC, activated B cell; GCB, germinal center B cell; MCL, mantle cell lymphoma; MZL, marginal zone lymphoma; CLL, chronic lymphocytic leukemia; PMBCL, primary mediastinal large B cell lymphoma; CTCL, cutaneous T cell lymphoma; ALCL, anaplastic large cell lymphoma; PTCL-NOS, peripheral T cell lymphoma-not otherwise specified; HL, Hodgkin lymphoma.

| HISTOLOGY | CELL LINE | LONCASTUXIMAB TESIRINE ($IC_{50}$, PM) | SG3199 ($IC_{50}$, PM) | RB4V1.2 NORMALIZED ANTIBODY BINDING CAPACITY | B12-C220-SG3249 ($IC_{50}$, PM) |
|---|---|---|---|---|---|
| ABC DLBCL | HBL-1 | 12.1 | 1.2 | 57597 | 2500 |
| ABC DLBCL | OCI-Ly-3 | 10 | 0.8 | 29163 | 1200 |
| ABC DLBCL | RCK8 | 35 | 1.2 | 88579 | 2500 |
| ABC DLBCL | RI-1 | 45 | 5.8 | 44683 | 3162.5 |
| ABC DLBCL | SU-DHL-2 | 400 | 1.1 | 10896 | 1300 |
| ABC DLBCL | TMD8 | 6 | 0.5 | 42808 | 550 |
| ABC DLBCL | U2932 | 1100 | 8.8 | 51776 | 5562.5 |
| GCB DLBCL | DB | 100 | 3.2 | 100060 | 300 |
| GCB DLBCL | DOHH2 | 0.5 | 0.5 | 15703 | 125 |
| GCB DLBCL | FARAGE | 2 | 0.9 | 132705 | 1650 |
| GCB DLBCL | KARPAS 422 | 31.1 | 5.3 | 147515 | 8000 |
| GCB DLBCL | OCI-Ly-1 | 1.4 | 1.4 | 17514 | 275 |
| GCB DLBCL | OCI-Ly-18 | 0.6 | 0.5 | 0 | 150 |
| GCB DLBCL | OCI-Ly-19 | 0.8 | 0.5 | 87842 | 175 |
| GCB DLBCL | OCI-Ly-7 | 1.4 | 1.1 | 68590 | 2250 |
| GCB DLBCL | OCI-Ly-8 | 0.3 | 0.5 | 58172 | 90 |
| GCB DLBCL | PFEIFFER | 790 | 14.6 | 77045 | 25000 |
| GCB DLBCL | SU-DHL-10 | 2.8 | 0.9 | 51975 | 750 |
| GCB DLBCL | SU-DHL-16 | 2812.5 | 1.1 | 25787 | 2500 |
| GCB DLBCL | SU-DHL-4 | 9.5 | 1.4 | 176138 | 2250 |
| GCB DLBCL | SU-DHL-5 | 2 | 0.8 | 56953 | 650 |
| GCB DLBCL | SU-DHL-6 | 6.3 | 14.6 | 63374 | 9500 |
| GCB DLBCL | SU-DHL-8 | 1.5 | 1.2 | 66635 | 2500 |
| GCB DLBCL | TOLEDO | 13 | 1.4 | 75346 | 2812.5 |
| GCB DLBCL | VAL | 0.4 | 0.5 | 74808 | 340 |
| GCB DLBCL | WSU-DLCL2 | 3 | 1.8 | 9848 | 1500 |
| PMBCL | KARPAS 1106P | 1.5 | 0.6 | 126819 | 690 |

| | | | | | |
|---|---|---|---|---|---|
| MCL | GRANTA519 | 1.4 | 0.5 | 50553 | 400 |
| MCL | JEKO1 | 5.3 | 0.5 | 45616 | 2250 |
| MCL | JVM2 | 5.5 | 2.0 | 47160 | 3000 |
| MCL | MAVER1 | 2.8 | 0.7 | 98854 | 900 |
| MCL | MINO | 1 | 0.5 | 115483 | 450 |
| MCL | REC1 | 20000 | 32.2 | 80734 | 32500 |
| MCL | SP49 | 1.3 | 0.5 | 40871 | 700 |
| MCL | SP53 | 2 | 0.5 | 72511 | 850 |
| MCL | UPN1 | 0.9 | 0.8 | 50600 | 600 |
| MCL | Z138 | 1.5 | 0.5 | 22049 | 300 |
| CLL | MEC1 | 5.5 | 0.5 | 92461 | 1800 |
| CLL | PCL-12 | 26 | 1.1 | 60314 | 2562.5 |
| MZL | ESKOL | 3 | 0.5 | 51311 | 650 |
| MZL | HAIR-M | 9.5 | 0.8 | 52925 | 900 |
| MZL | HC1 | 0.5 | 0.5 | 58951 | 200 |
| MZL | KARPAS 1718 | 0.7 | 0.5 | 36730 | 400 |
| MZL | SSK41 | 2 | 0.8 | 20233 | 650 |
| MZL | VL51 | 550 | 0.5 | 7497 | 450 |
| HL | AM-HLH | 600 | 0.8 | 0 | 300 |
| HL | KM-H2 | 2750 | 5.0 | 219 | 5250 |
| HL | L-428 | 14000 | 29.2 | 0 | 13750 |
| PTCL-NOS | FE-PD | 850 | 0.5 | 0 | 750 |
| ALCL | KARPAS 299 | 11500 | 17.5 | 0 | 12500 |
| ALCL | KI-JK | 4000 | 3.5 | 0 | 2962.5 |
| ALCL | L-82 | 5750 | 1.2 | 0 | 5500 |
| ALCL | SU-DHL-1 | 700 | 0.8 | 0 | 600 |
| CTCL | MAC1 | 900 | 0.5 | 0 | 1500 |
| CTCL | H9 | 1500 | 2.3 | 0 | 900 |
| CTCL | HH | 35000 | 23.4 | 0 | 24000 |
| CTCL | HUT-78 | 3500 | 0.8 | 0 | 1700 |
| Canine B cell lymphoma | CLBL1 | 175 | 0.5 | 0 | 175 |
| Murine B cell lymphoma | A20 | 2012.5 | 1.0 | 0 | 850 |
| Murine B cell lymphoma | BCL1 clone 5B1b | 500 | 0.5 | 0 | 435 |

**Table S3. $IC_{50}$ values obtained in DLBCL cell lines after 72 hours exposure to R-CHOP.**

| CELL LINE | R-CHOP (IC50, MG/ML) |
|---|---|
| DB | 0.0476 |
| DOHH2 | 0.0150 |
| FARAGE | 0.0009 |
| HBL-1 | 0.0586 |
| KARPAS 422 | 0.0325 |
| OCI-LY-1 | 0.0066 |
| OCI-LY-18 | 0.0529 |
| OCI-LY-19 | 0.0077 |
| OCI-LY-3 | 0.0139 |
| OCI-LY-7 | 0.0082 |
| OCI-LY-8 | 0.0005 |
| PFEIFFER | 0.1436 |
| RCK8 | 0.0431 |
| RI-1 | 0.8127 |
| SU-DHL-10 | 0.0010 |
| SU-DHL-16 | 0.1338 |
| SU-DHL-4 | 0.2383 |
| SU-DHL-5 | 0.0113 |
| SU-DHL-6 | 0.0134 |
| SU-DHL-8 | 0.0029 |
| SU-DHL-10 | 0.0409 |
| TMD8 | 0.0134 |
| TOLEDO | 0.0771 |
| U2932 | 0.1542 |
| VAL | 0.0003 |
| WSU-DLCL2 | 0.2056 |

**Table S3**. **Mutational analysis in B cell lymphoma cell lines.** Mapping based on Hg19 genome reference.

**Supplementary figures**

**Figure S1. Distribution of $IC_{50}$ values of loncastuximab tesirine (ADCT-402) between B and T derived lymphoma cell lines.** ****, P<0.0001 as determined by Mann-Whitney test.

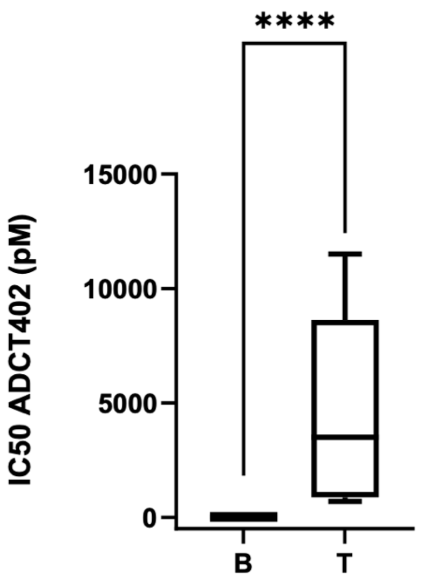


**Figure S2. Representative images of the apoptosis induction in two DLBCL cell lines exposed to loncastuximab tesirine (ADCT-402) and analyzed for annexin V by flow cytometry.** Cells were treated (2 x IC50) for 96 hours. The frequencies of annexin V positive cells (early apoptotic cells), annexin V/ propidium iodide double positive (late apoptosis), necrotic (annexin V negative, propidium iodide positive) and alive cells (annexin V/propidium iodide double negative) are shown.

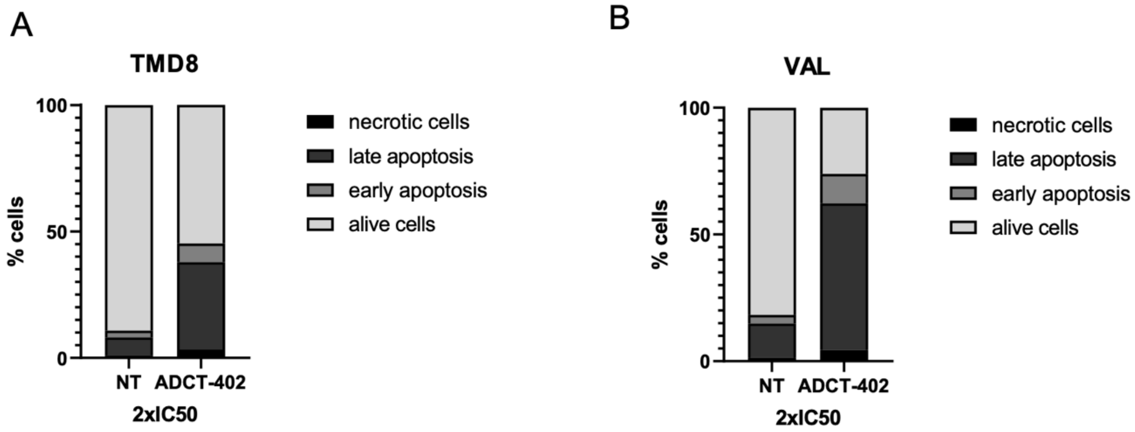

**Figure S3. Distribution of $IC_{50}$ values of loncastuximab tesirine among DLBCL cell lines based on the presence of absence of BCL2 and MYC chromosomal translocations, as single or concomitant events (double hit) and of TP53 status.** A) DLBCL cell lines with (n=16) and without (n=7) *TP53* inactivation. B) DLBCL cell lines with (n=15) and without (n=11) *BCL2* translocation. C) DLBCL cell lines with (n=7) and without (n=19) concomitant *BCL2* and *MYC* translocation. D) DLBCL cell lines with (n=10) and without (n=16) *MYC* translocation. *, P< 0.05; **, P< 0.001.

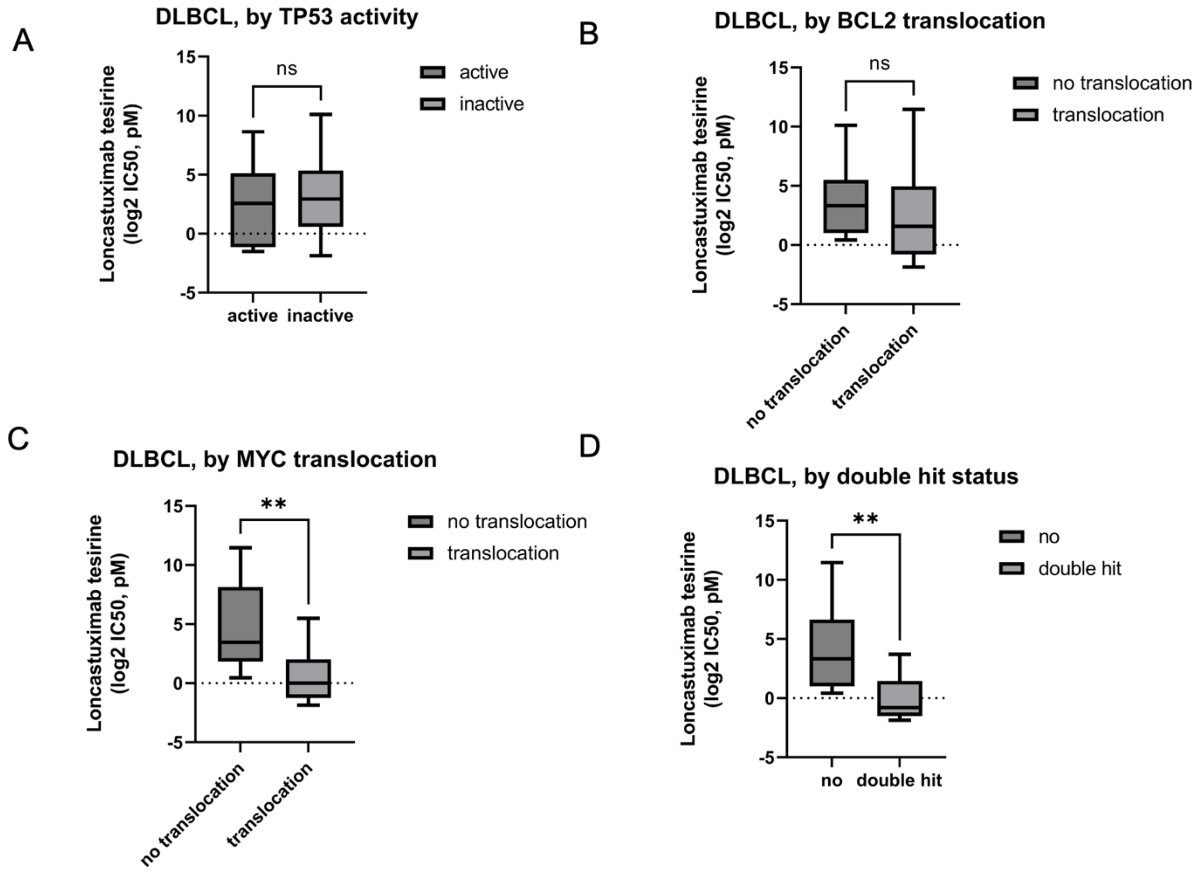

**Figure S4. MYC, CD19 and $IC_{50}$ values of loncastuximab tesirine among DLBCL cell lines.** A) CD19 surface expression between cell lines with and without MYC translocation. B) CD19 RNA expression values measured via RNA-Seq between cell lines with and without MYC translocation. C) CD19 RNA expression values measured via microarray between cell lines with and without MYC translocation. D) correlation plot between CD19 surface expression and MYC expression measured via RNA-Seq. E) correlation plot between CD19 RNA and MYC expression, both measured via RNA-Seq. F) Correlation between loncastuximab tesirine $IC_{50}$ values and MYC expression measured via RNA-Seq.

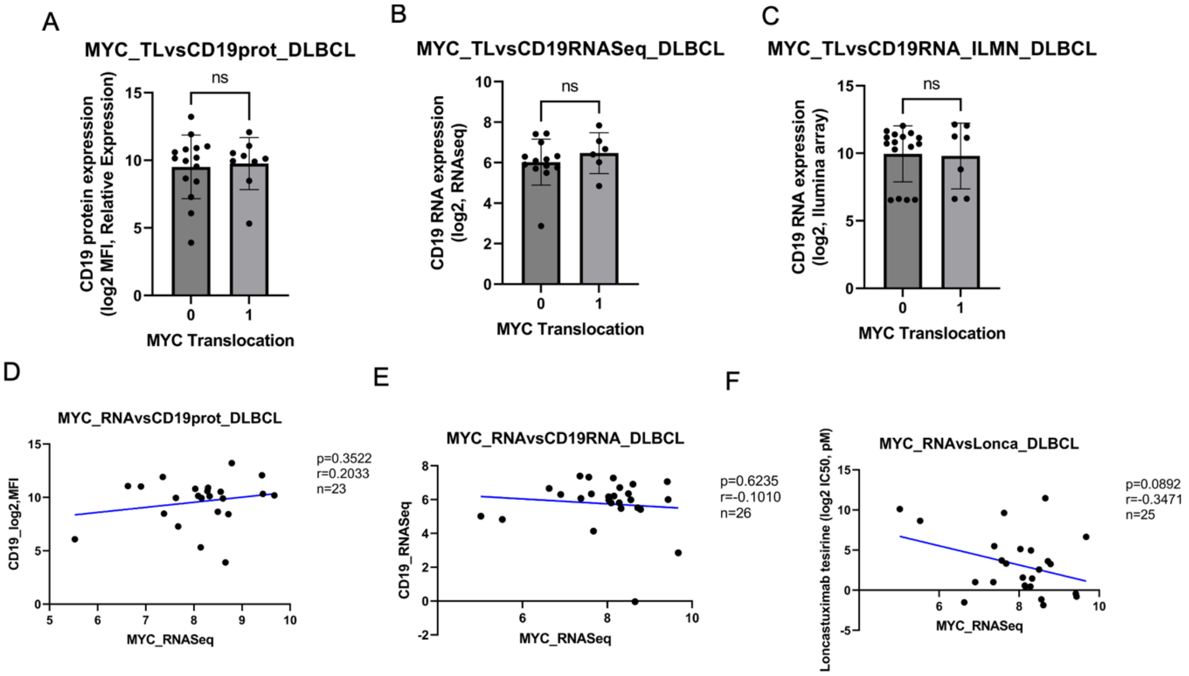


**Figure S5. Distribution of $IC_{50}$ values of SG3199 between B and T derived lymphoma cell lines.**

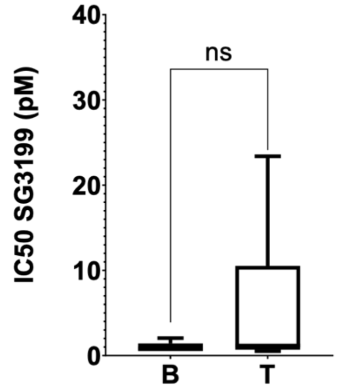

**Figure S6. Pearson correlation between SG3199 and CD19 absolute expression among B cell lymphoma cell lines.**

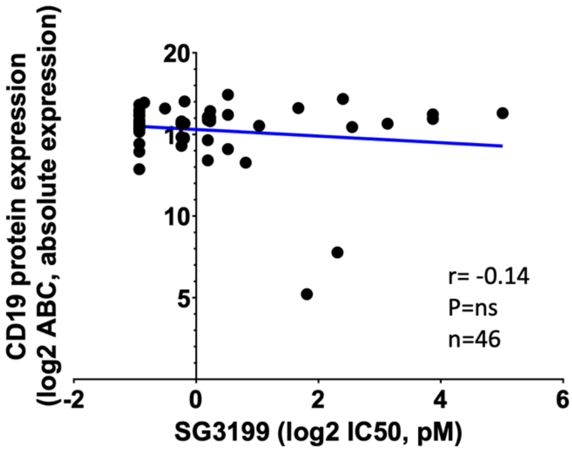


**Figure S7. Distribution of $IC_{50}$ values of loncastuximab tesirine and SG3199 across all cell lines (A) and among B cell lymphoma cell lines (B).** ****, P<0.0001 as determined by Mann-Whitney test.

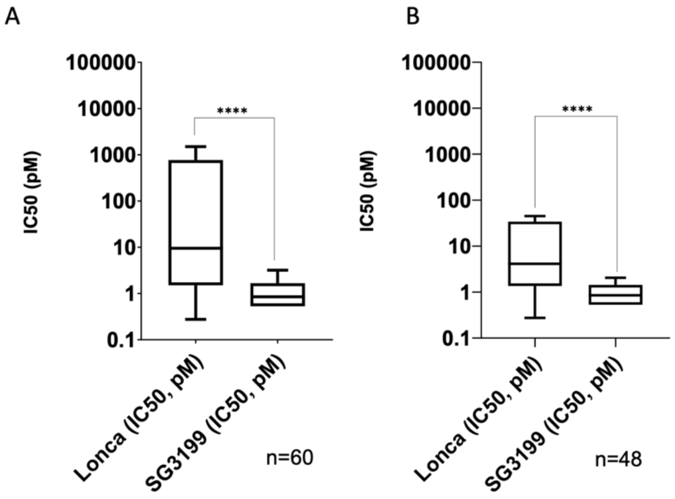

**Figure S8. Distribution of IC$_{50}$ values of SG3199 among DLBCL cell lines based on the presence of absence of BCL2 and MYC chromosomal translocations, as single or concomitant events (double hit) and of TP53 status.** A) DLBCL cell lines with (n=16) and without (n=7) *TP53* inactivation. B) DLBCL cell lines with (n=15) and without (n=11) *BCL2* translocation. C) DLBCL cell lines with (n=7) and without (n=19) concomitant *BCL2* and *MYC* translocation. D) DLBCL cell lines with (n=10) and without (n=16) *MYC* translocation. *, P< 0.05; **, P< 0.001.

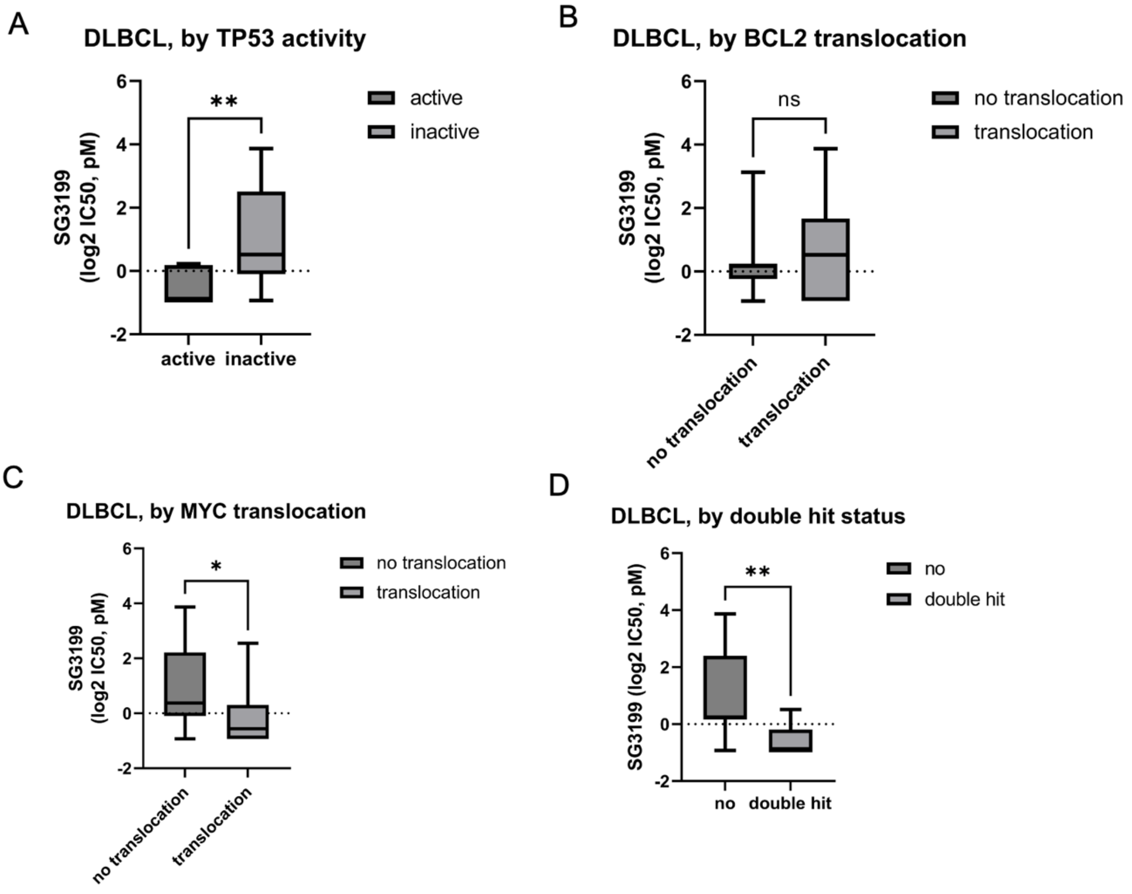

**Figure S9. Pearson correlation between SG3199 $IC_{50}$ values and MYC RNA levels measured** via RNA-Seq (A) **and** via microarray (B) **among DLBCL cell lines.**

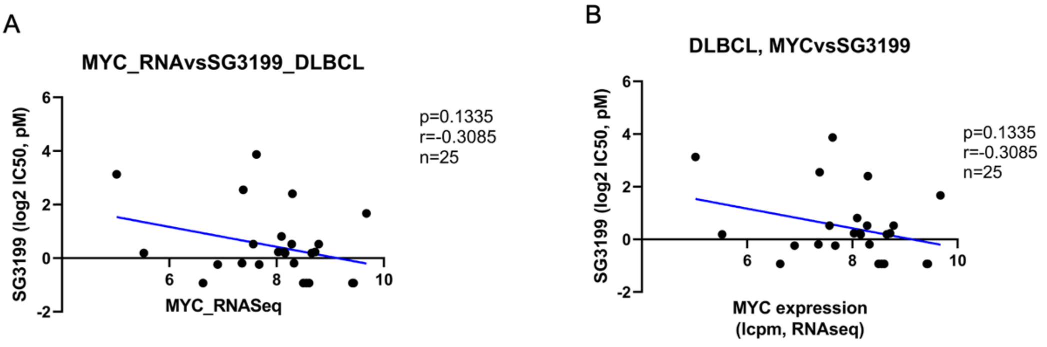


**Figure S10. SU-DHL-6 cell line is sensitive to loncastuximab tesirine and to its naked antibody, but it is resistant to its warhead SG3199.** Loncastuximab tesirine, SG3199 and rB4v1.2 activity was evaluated by MTT assay for 96 hours of treatment. X axis, concentration in pM; Y axis, fold to untreated. * q values < 0.05 of loncastuximab tesirine vs all other treatments (SG3199, rB4v1.2) was determined by unpaired t-test followed by two-stage step-up (Benjamini, Krieger, and Yekutieli) multiple comparisons, FDR(q)=0.05.

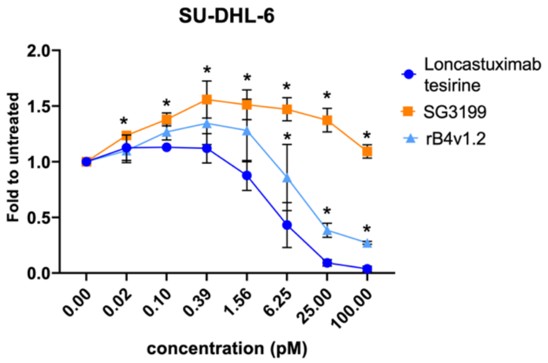


**Figure S11.** Correlation between **in vitro anti-proliferative activities of loncastuximab tesirine and two others anti-CD19 ADCs.** Pearson correlations between log2 $IC_{50}$ (pM) of loncastuximab tesirine activity with coltuximab ravtansine (SAR34199) (A) or huB4-DGN462 (B).

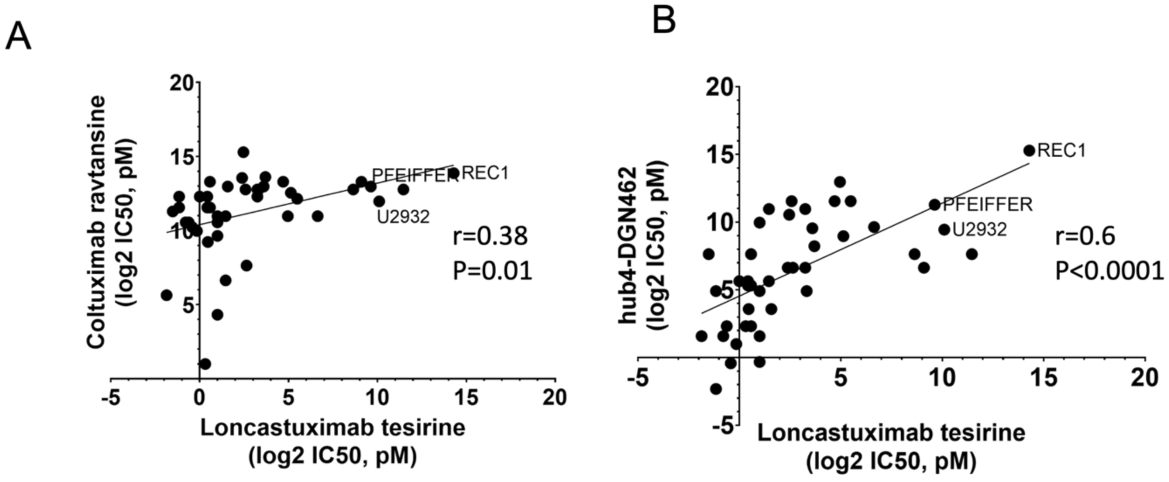

**Figure S12. Cell cycle distribution of DLBCL cell lines after 96h of treatment with loncastuximab tesirine alone or in combination with venetoclax, idelalisib or copanlisib.** TMD8 (A), WSU-DLCL2 (B), OCI-LY-3 (C) and VAL (D) were treated with two times $IC_{50}$. Statistics were calculated with the Student's t-test. *P value < .05. Single asterisk compares exposure conditions with the untreated sample.

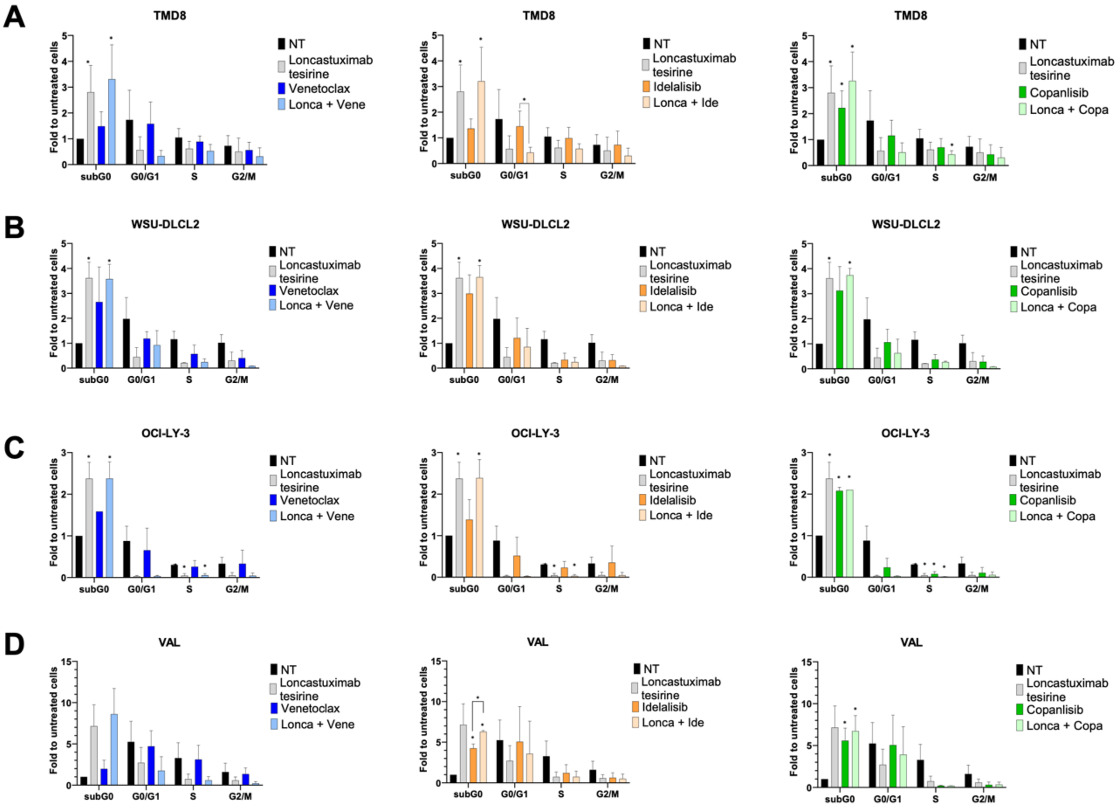

**Figure S13. Quantification of protein changes in cells treated with loncastuximab tesirine as single agent or in combination with venetoclax, idelalisib and copanlisib.**

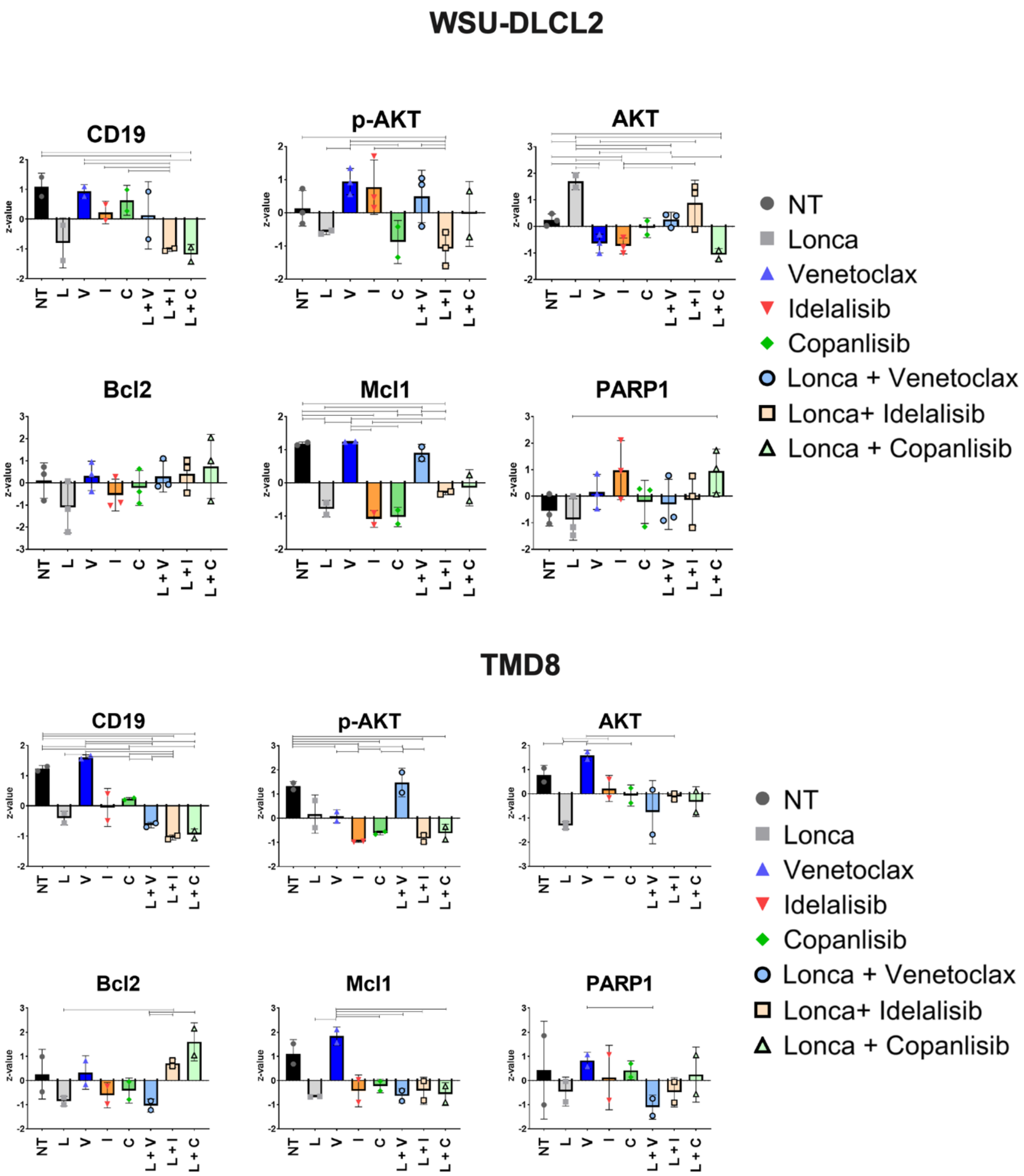

**Figure S14. Single agent in vivo activity of loncastuximab tesirine and B12-SG3249 in TMD8 ABC DLBCL.**

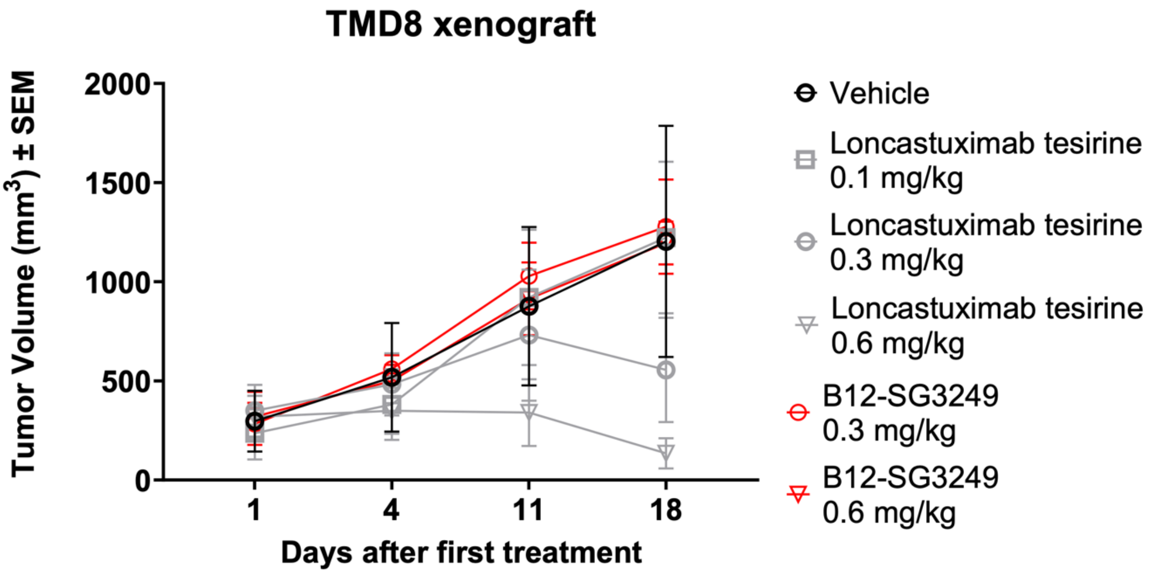


**Figure S15. Single agent in vivo activity of copanlisib in TMD8 ABC DLBCL.** * p values < 0.05 of copanlisib (both dosages) vs vehicle, as determined by Mann-Whitney test.

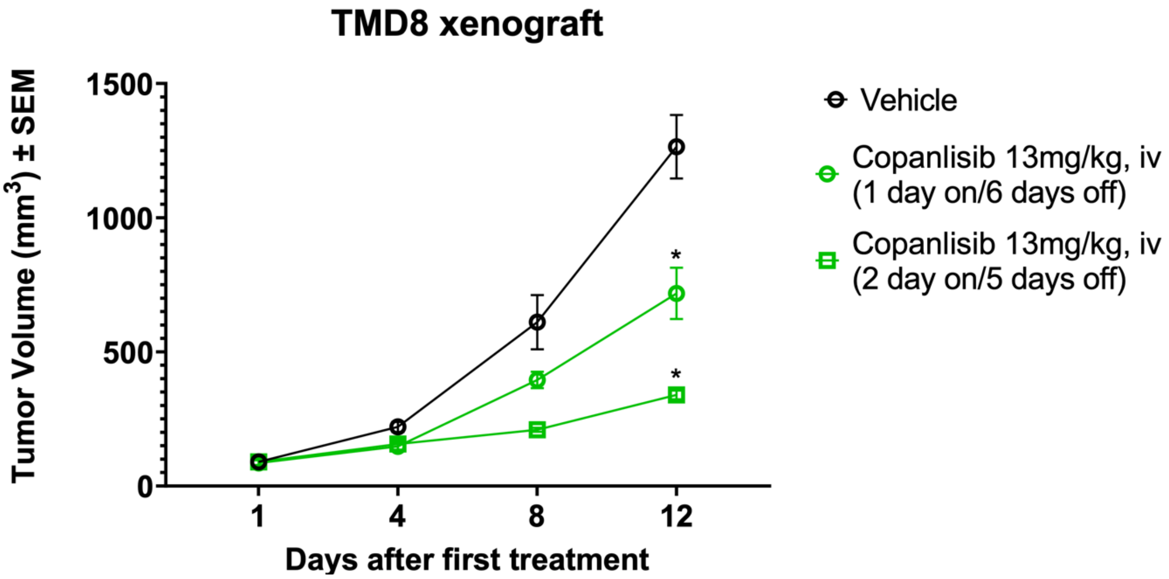

**Figure S16. Tumor weight in grams of xenografts from mice treated with loncastuximab tesirine in combination with copanlisib (A) or venetoclax (B).** *, P<0.05; **, P<0.005; ***, P<0.0001 as determined by Mann-Whitney test.

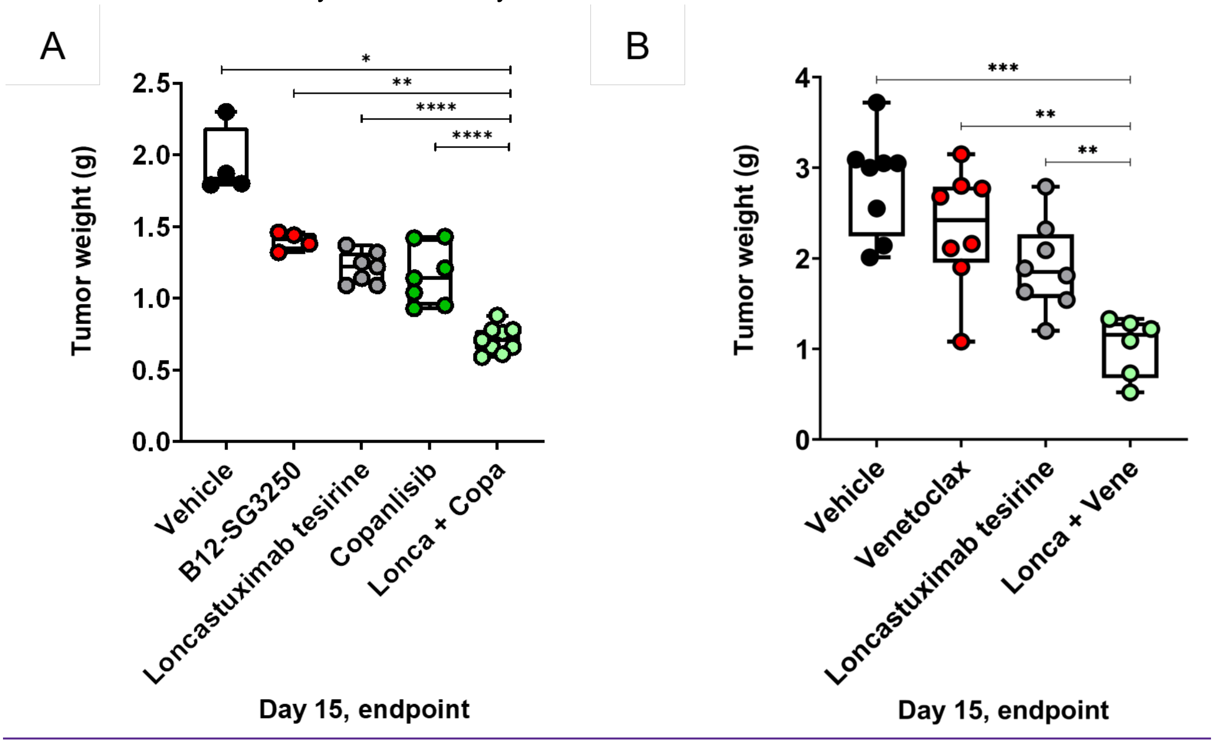
